# Diet-induced chromatin states influence intestinal stem cell memory

**DOI:** 10.64898/2026.02.19.706894

**Authors:** Dominic R Saiz, Yesenia Barrera Millan, Thomas Hartley McDermott, Gabriella Cerna, Swathi Sankar, Fiona Farnsworth, Eric Uher, Gourab Lahiri, Kelly Lintecum, Karla Mullen, Benjamin B Bartelle, Miyeko D Mana

**Author notes:** CONTACT INFORMATION: Miyeko D. Mana.

## Abstract

Intestinal stem cells (ISCs) integrate dietary cues through a metabolic–transcriptional axis, but whether these mechanisms create a lasting epigenetic memory remains unclear. Here, we investigate diet-induced chromatin adaptations in ISCs using a high-fat Western diet (HFD) mouse model. HFD broadly remodels chromatin accessibility, altering pre-existing open regulatory regions. Many differentially accessible regions (DARs) persist during the differentiation of ISCs into transient amplifying cells (TACs). Notably, HFD-induced DARs are retained following diet normalization despite phenotypic reversibility, and HFD re-exposure enhances ISC self-renewal and adenoma growth compared with naïve HFD exposure. The HFD-induced chromatin changes require *Ppar-d/a* nuclear receptors but are independent of their transcriptional targets. Although a subset of HFD-induced DARs is maintained following *Apc* loss, extensive chromatin remodeling driven by tumor suppressor inactivation largely overrides diet-dependent differences in *Lgr5*⁺ ISCs. Together, these findings demonstrate that ISCs retain a chromatin-based memory of dietary fat exposure.

## INTRODUCTION

Adult stem cells maintain tissue homeostasis and enable regeneration in health and disease (Morrison and Spradling, 2008; van der Flier and Clevers, 2009). They sense and respond to environmental cues for regulation and determination of cell fate. Diet plays a pivotal role in stem cell behavior by modulating nutrient availability (Jackson and Finley, 2024; Meacham et al., 2022). Nutritional inputs directly alter the metabolic state and transcriptional control, serving as drivers of stemness, proliferation and differentiation (Mihaylova et al., 2014; Novak et al., 2021). Less clear is whether stem cells adapt to diet by altering their chromatin state and establishing a molecular memory.

Intestinal stem cells (ISCs) lie at the base of the intestinal crypt surrounded by a supportive microenvironment that facilitates a differentiation program that spurs constant upward migration of absorptive and secretory cells (Barker et al., 2007). ISCs respond to nutrient cues through diverse strategies. For example, caloric restriction invokes ISC self-renewal through non-cell autonomous signals from niche Paneth cells (Yilmaz et al., 2012). Ketogenic conditions induce β-hydroxybutyrate (βOHB) production via HMGCS2 that modulates nuclear Notch activity in ISCs (Cheng et al., 2019). We recently showed that a high-fat Western diet (HFD) relies on a PPARd/a nuclear receptor program and the downstream transcriptional target, *Cpt1a*, to control the mitochondrial oxidation of long-chain fatty acids (Beyaz et al., 2021b, 2016; Mana et al., 2021). Loss of *Ppar-d/a*, or separately *Cpt1a*, abrogates the HFD-induced increase in stem cell function, proliferation, and regenerative capacity, linking a metabolic-transcriptional axis to stemness and tumorigenic potential in this dense nutrient state (Mana et al., 2021). Conversely, a 24hr nutrient-restricted fast similarly induces a PPAR-CPT1a program to maintain the ISC pool and coordinate homeostasis (Mihaylova et al., 2018).

While dietary interventions have been used to dissect the metabolic-transcriptional axis, how these diets reshape chromatin landscapes and contribute to longer-term cellular memory is an emerging frontier. The fasted state stimulates the production of βOHB leading to a prominent enrichment of H3K9bhb on regulatory elements of metabolic genes in crypts, integrating nutrient availability with chromatin remodeling (Terranova et al., 2021). A HFD induces DNA methylation changes in the colonic epithelium with many of the differentially methylated regions persisting transiently after switching mice to a low-fat control diet, underscoring diet-driven epigenetic memory at the tissue level (Li et al., 2018). How ISC chromatin alterations facilitate the adaptation to dietary exposure is less characterized. Using a pro-obesity HFD driving altered ISC programs and increased risk of transformation (Beyaz et al., 2021b, 2016; Mana et al., 2021), we investigate the ISC chromatin accessibility and test the ability of these tissue specific stem cells to retain memory of prolonged lipid exposure.

## RESULTS

### HFD Alters Chromatin Accessibility in ISCs and TAC progenitors

Diet alters nutrient availability for metabolic signaling and promotes transcriptional changes that influence ISC behavior. As demonstrated previously, a high-fat Western diet (HFD) induces a phenotype characterized by an increase in ISC numbers, proliferation, self-renewal, and capacity to generate adenomas (Beyaz et al., 2016; DeClercq et al., 2015; Mah et al., 2014; Mana et al., 2021). These changes are associated with an increase of multiple histone modifications in HFD-fed crypt cells relative to control diet (CD) counterparts as seen by Western blot (Figures S1A and S1B), highlighting the availability of metabolic substrates for histone labeling. To gain further insights into the chromatin changes, we began by assessing the chromatin landscape using ATAC-seq (Buenrostro et al., 2015) on small intestinal ISC and transient amplifying cell (TAC) populations (*Lgr5*^GFPhi^ ISCs and *Lgr5*^GFPlow^ progenitors, respectively) isolated from mice (*Lgr5*^eGFP-IRES-CreER^) (Barker et al., 2007) on a six-month CD or HFD (Figure S1C).

Broad comparison across all four populations reveal marked differences in chromatin accessibility between ISCs and TACs (Figure 1A). ISCs show clear diet-dependent accessibility shifts, whereas the TAC populations remain distinct from ISCs but display reduced diet-induced accessibility changes. Within ISCs, the HFD induces more open chromatin than the CD, observed by categorizing the top 10,000 open genomic regions (Figure S1D). However, neither diet biases more closure of the chromatin, indicating that gains in accessibility are more pronounced in the HFD (Figure S1D).

**Figure 1.**
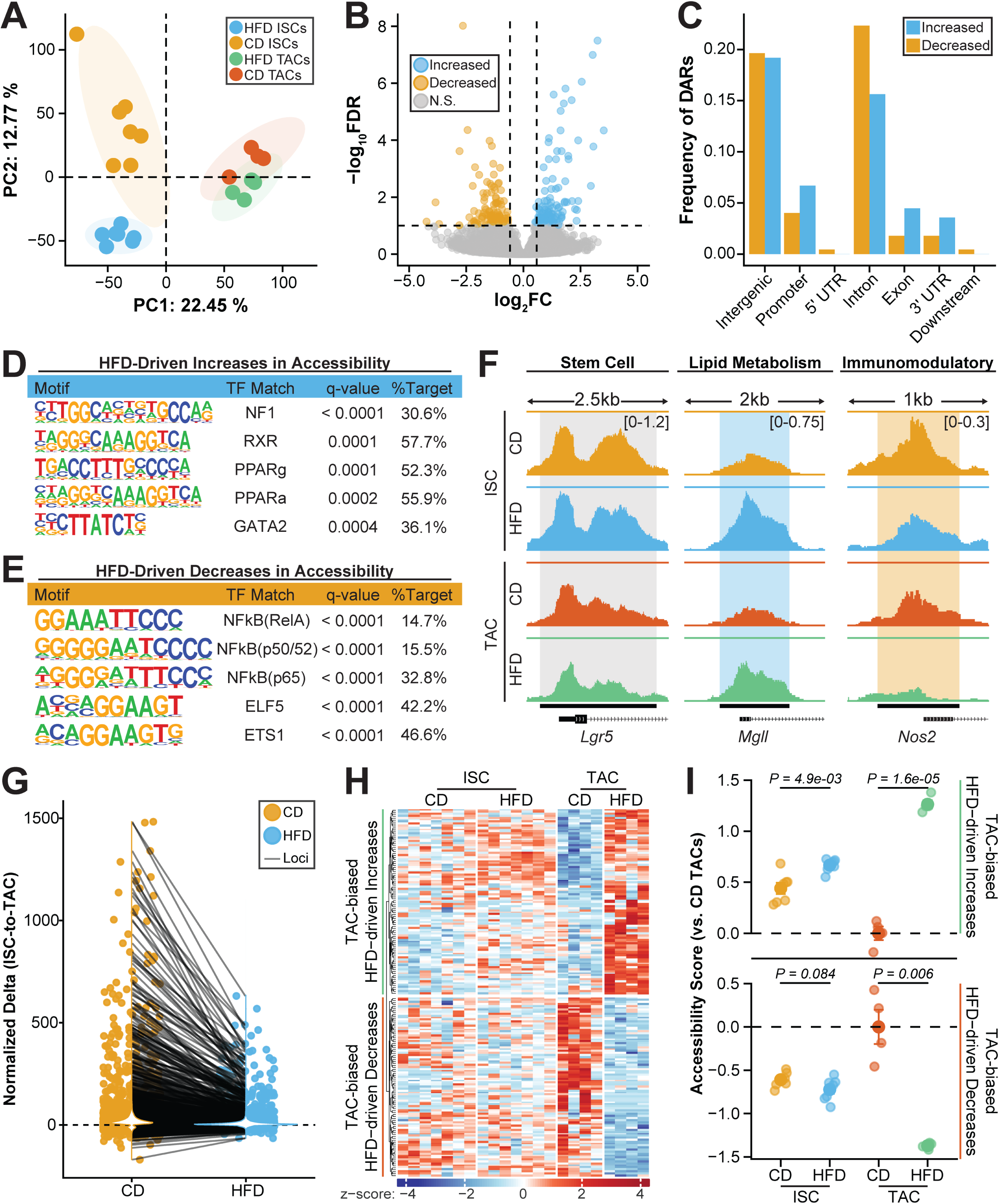
HFD alters the chromatin accessibility landscape in ISCs and TACs. **(A)** Principal component analysis (PCA) plot of variance-stabilized chromatin accessibility values (counts) for ISCs and TACs under CD or HFD conditions. Each point represents one mouse-derived ATAC-seq sample. WT ISCs: CD n=7, HFD n=7; WT TACs: CD n=4, HFD n=4. **(B)** Volcano plot showing DARs between CD and HFD-exposed ISCs. Each point represents a genomic region: blue = increased accessibility, orange = decreased accessibility, and gray = nonsignificant. Significance threshold: false discovery rate (FDR) < 0.1; |log_2_FC| > log₂ 1.5. **(C)** Bar plot showing the frequency of increased and decreased DARs across genomic annotation categories. **(D,E)** Motif-enrichment analysis for HFD-increased **(D)** and HFD-decreased **(E)** regions displaying the ranked enriched transcription factor motifs, corresponding transcription factor (TF) matches, q-values, and percent of target loci containing the matching TF motif. **(F)** Representative genome-browser tracks of chromatin accessibility at the *Lgr5*, *Mgll*, and *Nos2* loci in ISCs and TACs under CD and HFD. Shaded areas denote differential peak region. **(G)** Paired scatter plot of normalized chromatin accessibility differences between loci exhibiting significant ISC-to-TAC accessibility changes (ΔISC–TAC) in CD (orange) and baseline-corrected HFD (blue) conditions. 5,889 regions show attenuation of Δ ISC–TAC magnitude in HFD (>50% reduction), including 3,863 regions with 50–75% attenuation and 2,023 regions with >75% attenuation. Wilcoxon signed rank test *P < 2.2e-16*. **(H)** Heatmap of z-scored accessibility values for TAC-exclusive HFD-increased and HFD-decreased regions in ISCs and TACs under CD and HFD. **(I)** Dot plots displaying accessibility module scores for HFD-increased and HFD-decreased regions in ISCs and TACs, with *P*-values from two-tailed unpaired *t*-tests.

To identify the significant changes in the ISC chromatin landscape induced by the HFD, we analyzed differentially accessible regions (DARs: genomic loci with significant differences in number of reads (False Discovery Rate [FDR] < 0.1, |log2FoldChange| > log2(1.5)). We identified similar number of DARs between CD and HFD ISCs (Figure 1B). The DARs are located throughout genomic elements and concentrate in distal intergenic regions and introns (Figure 1C), suggesting regulation outside direct promoter-driven transcription. Overall, DARs exhibit little diet-induced bias towards specific genomic regional annotations as similar number of regions gained and lost accessibility. However, the promoters represented in the DARs (TSS±2kb) are differentially enriched in transcription factor binding sites that infer diet-induced regulatory programs, consistent with known transcriptional activity (Beyaz et al., 2016; Mana et al., 2021) (Figure S1G). For example, DARs open in the HFD are enriched with PPAR and RXR binding sites (Figure 1D). Similarly, gene set enrichment analysis and molecular pathways associated with these HFD DARs are enriched in abundant metabolic programs with fatty acid metabolism and PPAR signaling prominently represented (Figures S1E and S1F). These are consistent with known roles in mediating the HFD-induced ISC phenotypic signature (Beyaz et al., 2016; Mana et al., 2021). DARs open in the CD, and therefore more closed in the HFD, are enriched with NFkB binding sites (Figure 1E). In general, the DARs reflect a difference of enhanced peaks rather than the opening of novel accessible regions, indicating that the HFD acts on existing chromatin architecture to modulate regulation (Leung et al., 2016; Siersbæk et al., 2017). Accessibility peaks can be visualized using representative peaks and statistically validated at loci attributed to lipid metabolism (*Mgll, Fabp1*, *Acaa1b*, *Cpt2*), and immune modulation (*Nos2*, *Gbp10*, *Mlrp9b, Mir29b-2*) (Figure 1F and S1H). However, loci involved in stemness (*Lgr5*, *Olfm4*, *Ascl2, Axin2*) are not directly impacted by dietary exposure to the HFD, highlighting that diet-induced cellular metabolism and intracellular signaling are altered independently of factors maintaining ISC identity.

The HFD induces lipid metabolic programs that are shared by a fasted state, underscoring common molecular features of the dietary regimens (Mana et al., 2021; Mihaylova et al., 2018; Terranova et al., 2021). We were curious if these two disparate dietary states invoke similar chromatin changes. Indeed, a 24 hour fast induced rapid chromatin alterations in ISCs such that they share many HFD-induced DARs (Figure S2A). Heatmaps of HFD-increased and HFD-decreased peaks show that fasting induced accessibility changes seen with HFD, in which the number of fasted ISC peaks appear intermediate between CD and HFD. We calculated a module score for predefined peak sets (HFD-increased or HFD-decreased DARs), defined as the mean accessibility across all peaks in a given module for each sample. Consistently, module scores for HFD-increased peaks were significantly elevated in both HFD and fasted ISCs relative to CD, whereas scores for HFD-decreased peaks were significantly reduced in the fasted condition, with ISCs yielding values more similar to CD (Figure S2B), suggesting the ISCs are more poised to increase accessibility in response to this dietary program. Notably, the fasted state shows evidence of sex-specific differences in ISC chromatin accessibility that are not observed in the CD and HFD conditions. These differences introduced variation in the fasted DARs and reduced the significance of peak accessibility. More broadly examining accessibility throughout all samples, the fasted state is more similar to CD than HFD, with the fasted sex differences separating the samples more disparately (Figure S2C).

ISCs give rise to TACs over a mitotic division. To explore the nature of this transition in CD and HFD, we first examined the difference of trajectory in a HFD condition normalized to that of the CD condition. We find that comparing individual peaks in this transition reveals reduced shifts in accessibility between the cell states in a HFD (Figure 1G), with the majority of the analyzed genomic loci having attenuated changes in the HFD ISC-to-TAC transition compared to the control counterpart, indicative that the HFD-induced effects persist within the lineage to retain a residual chromatin profile established in ISCs. To test if the ISC HFD peaks remain accessible in the TAC population, we assessed the trajectory of the differentially accessible variable regions along the ISC to TAC transition. In general, we identified discrete modalities of altered chromatin accessibility upon differentiation that can be classified into three groups. The first group is largely consistent with transcriptional programs that follow canonical differentiation patterns and are not influenced by the HFD (Figure S3A). We find that peaks in ISCs either lose accessibility in progenitors, such as those related to stemness specific Wnt and Hippo signaling, or gain accessibility in the progenitors, such as peaks linked to the PPAR and retinol metabolism pathways whose activity increase with, oxidative metabolism upon differentiation (Figure S3B). Transcriptional profiles were compared to the accessibility differences within this modality group, revealing a significant correlation of changes in accessibility near the TSS to changes in transcriptional outcomes (Figure S3C). The remaining TAC DARs exhibit two divergent differentiation patterns in the HFD condition (Figure S3D) both sharing common KEGG enrichment features of ISCs, such as PPAR signaling (Figure S3E) and a significant correlation of transcriptional differences in TACs (Figure S3F). This creates a second modality in which peaks are maintained along the ISC to progenitor cell divisions, such that DARs established in ISCs are maintained in the progenitors, allowing for the continuation of the HFD phenotype and expansion of stem-like cells (Figures S3G and S3H).The third modality identifies additional DARs exclusive to the HFD-exposed progenitors (Figures 1H and 1I). These distinct regions are similarly enriched with lipid metabolism mediators (PPAR, RXR and HNF4a) in open peaks (Figure S3I), and immune modulating NFkB/RelA in HFD-induced closed peaks (Figure S3J), thereby adding more features observed in ISCs to new progenitor pools, and extending the metabolic and immune-modulatory hallmark of dietary exposure along the intestinal differentiation hierarchy.

### *Ppar-d/a* lipid metabolism is required for HFD-induced chromatin changes

PPARd/a are major lipid-activated nuclear receptors mediating the adaptation to a HFD(Mana et al., 2021). Genetic loss of intestinal *Ppar-d/a* (*Ppar-d/a*^iKO^: *Ppard*^fl/fl^; *Ppara*^-/-^; *Villin*^CreER^; *Lgr5*^eGFP-IRES-CreER^) abrogates the ISC HFD phenotype, thereby reducing proliferation and regenerative capacity without affecting long-term ISC maintenance as observed by the longevity of the double mutants maintained on a CD or HFD regimen(Mana et al., 2021). Given that PPAR binding sites are prominent in differentially accessible peaks induced by the HFD, we tested how these transcription factors influence the chromatin landscape. We conducted ATAC-seq on isolated *Lgr5*^GFPhi^ ISCs from *Ppar-d/a*^iKO^ mice maintained on a CD or HFD from 8 weeks of age (Figure 2A). Irrespective of diet, in the PCA *Ppar-d/a*^iKO^ samples cluster between the wildtype (WT) CD and HFD samples, suggesting similarity within *Ppar-d/a*^iKO^ ISCs and neutralization of accessible peaks in the diet conditions (Figures 2B and 2C). Indeed, the significant DARs identified in WT HFD ISCs are substantially diminished in absence of the PPARs under either dietary regimen (Figure 2D and 2E). Promoter DARs related to genes involved in lipid metabolism are reduced to near CD levels (Figure 2F). Similarly, the significant HFD-driven decreased DARs are also reduced with loss of intestinal *Ppar-d/a* (Figure 2E), further observing that the immune-associated genes with elevated peaks in regulatory regions under the control diet are diminished in the *Ppar-d/a*^iKO^ ISCs (Figures 2G and S4A). In contrast, peaks in ISC marker genes such as *Lgr5*, *Olfm4* and *Ascl2* remain unaffected, demonstrating the independence of the stemness program (Figure S4A). We performed a pairwise comparison of the significant DARs found in *Ppar-d/a*^iKO^ which revealed only three HFD-driven DARs in the double knockout ISCs (Figure2C). This number drops further when comparing the DARs between *Ppar-d/a*^iKO^ and WT (Figure S4B). These data suggest that the PPARs are required for the HFD-induced chromatin changes and highlight how essential the PPARs are for responding and adapting to the HFD.

**Figure 2.**
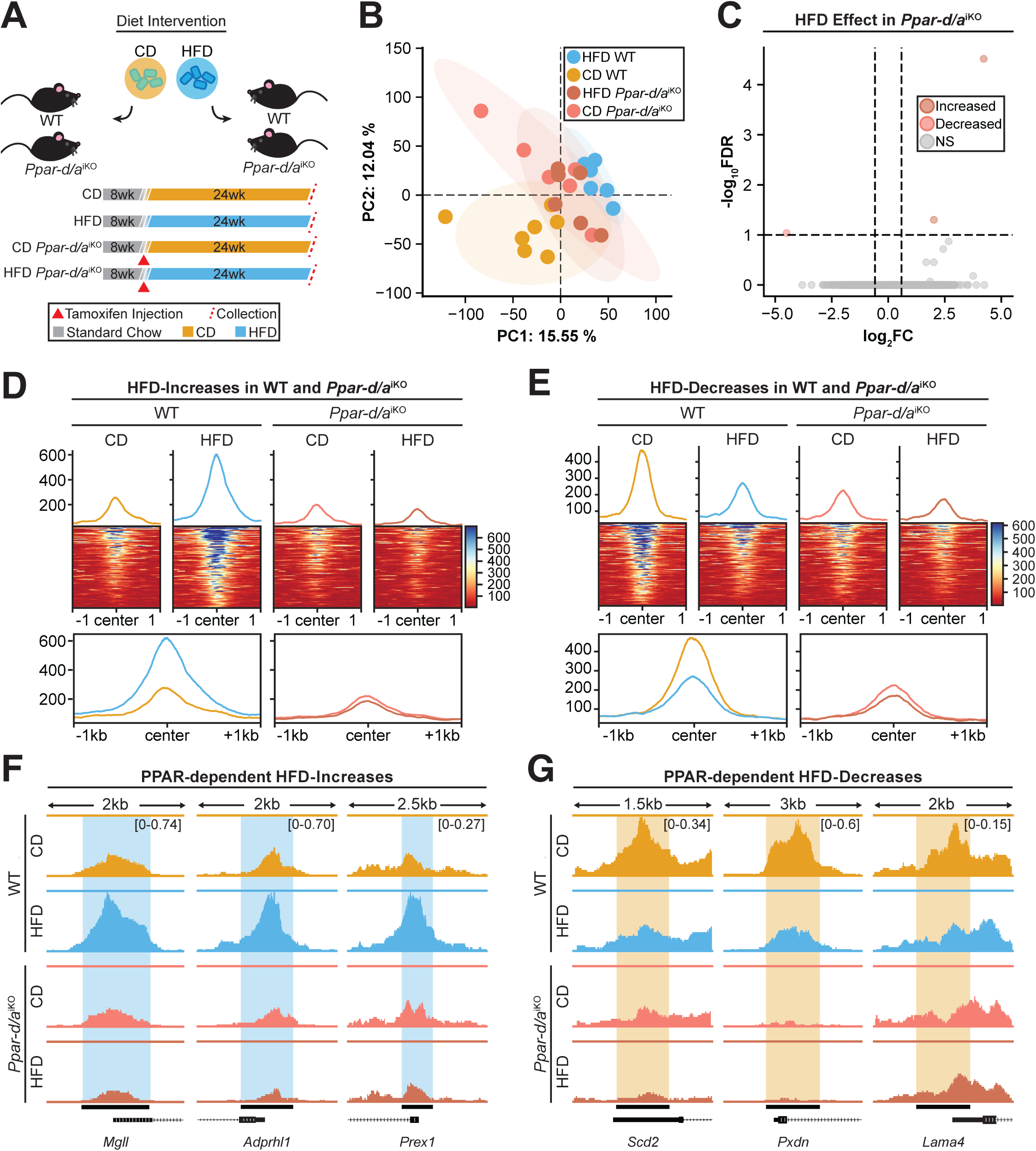
PPAR signaling influences HFD-induced chromatin accessibility changes in ISCs. **(A)** Experimental schematic depicting the dietary intervention and genetic models. *Ppar-d/a* intestinal knockout and WT mice were maintained on CD or HFD for 24 weeks prior to ISC isolation and analysis. **(B)** PCA plot of variance-stabilized chromatin accessibility profiles of ISCs from CD and HFD-fed WT and *Ppar-d/a*^iKO^ mice. Each point represents one mouse-derived ATAC-seq sample. WT ISCs: CD n=7, HFD n=7; *Ppar-d/a*^iKO^ ISCs: CD n=6, HFD n=6. **(C)** Volcano plot showing DARs in HFD *Ppar-d/a*^iKO^ compared with HFD WT ISCs. Each point represents a genomic region: orange = increased accessibility in HFD, red = decreased accessibility, and gray = nonsignificant. Significance threshold: FDR < 0.1; |log_2_fold-change| > log_2_1.5. **(D,E)** Average signal profiles and heatmaps showing accessibility changes centered around DARs increased **(D)** and decreased **(E)** under HFD in WT mice. Tracks display accessibility for CD WT, HFD WT, CD *Ppar-d/a*^iKO^, and HFD *Ppar-d/a*^iKO^. **(F,G)** Genome-browser tracks displaying chromatin accessibility with an **(F)** increase (*Mgll*, *Adprhl1*, *Prex1*) or **(G)** decrease (*Scd2*, *Pxdn*, *Lama4*) in HFD conditions. Enhanced peaks are *Ppar-d/a* dependent. Tracks are aligned to the transcription start site (TSS) with scale bars showing distance (kb) upstream and downstream.

PPAR activation rapidly rewires cellular metabolism, invoking a transcriptional response leading to the increased lipid trafficking and utilization. A key target, *Cpt1a*, is the rate-limiting enzyme for mitochondrial long-chain fatty acid oxidation (FAO). We previously determined the requirement of *Cpt1a* in facilitating the HFD-induced ISC phenotype (Mana et al., 2021). To test the ability of mitochondrial fatty acid metabolism to influence chromatin accessibility relative to the PPAR transcription factors, we performed ATAC-seq on *Cpt1a*-null ISCs (*Cpt1a*^iKO^: *Cpt1a*^fl/fl^; *Villin*^CreER^; *Lgr5*^eGFP-IRES-CreER^) from mice on a CD or HFD. DARs identified in WT ISCs were compared to the knockout ISCs. In the HFD-induced open state, *Cpt1a*^iKO^ ISC DARs are highly similar to the WT HFD ISCs (Figures S4C and S4D), suggesting minimal influence on the chromatin architecture with the loss of CPT1a-mediated FAO. This indicates that despite loss of mitochondrial FAO metabolites, chromatin accessibility under HFD conditions remains largely unchanged. These findings suggest that metabolic perturbations are buffered at the chromatin level, and accessibility is primarily reshaped by when transcriptional regulators that orchestrate the HFD response are disrupted.

### HFD induces a persistent chromatin state

Current evidence suggests that ISCs maintain a molecular memory of the HFD: first, multiple ISC DARs are carried into the progenitor population (Figure S3G and S3H), and second, the HFD-induced ISC regenerative capacity persists after 1 week CD normalization to allow turnover of the intestinal cells, although the HFD ISC functional clonogenic capacity is dissipated by 4 weeks (Beyaz et al., 2016). We continued to explore the extent that ISCs maintain features of the HFD after dietary reversion (Figure 3A). Phenotypically, we find that the ISC marker OLFM4 begins to identify fewer ISCs per crypt after dietary normalization but numbers remain elevated above control (Figures 3C and 3E). Additionally, ISC proliferation reduces rapidly upon removal of the HFD diet such that a 4-hour pulse of BrdU nucleoside analog no longer exhibits increased BrdU^+^ ISCs after 1 and 4 weeks HFD removal (Figures 3D and 3F). Molecularly, we see that protein levels of PPARd/a transcriptional targets, CPT1a and HMGCS2, are maintained in crypts after one week but the abundance reduces to control levels after 4-week removal from the HFD (Figure3B). Lastly, we tested the tumorigenic capacity of the ISCs by removing functional tumor suppressor *Apc* specifically in *Lgr5*^+^ stem cells (*Apc*^KO^: *Apc*^fl/fl^; *Lgr5*^eGFP-IRES-CreERT2^) to induce adenoma formation. We find that adenomatous lesions remain larger than the CD counterparts after 1-week removal from the diet, much like the HFD (Figures 3G and 3H). However, the growth of adenomas slows after 4-week normalization, resembling similar β-catenin^+^ lesion size as the CD adenomas, thereby losing the impact of the HFD (Figures 3G and 3H). Hence, after 4-week removal from the HFD, ISCs phenotypically resemble their CD counterparts.

**Figure 3.**
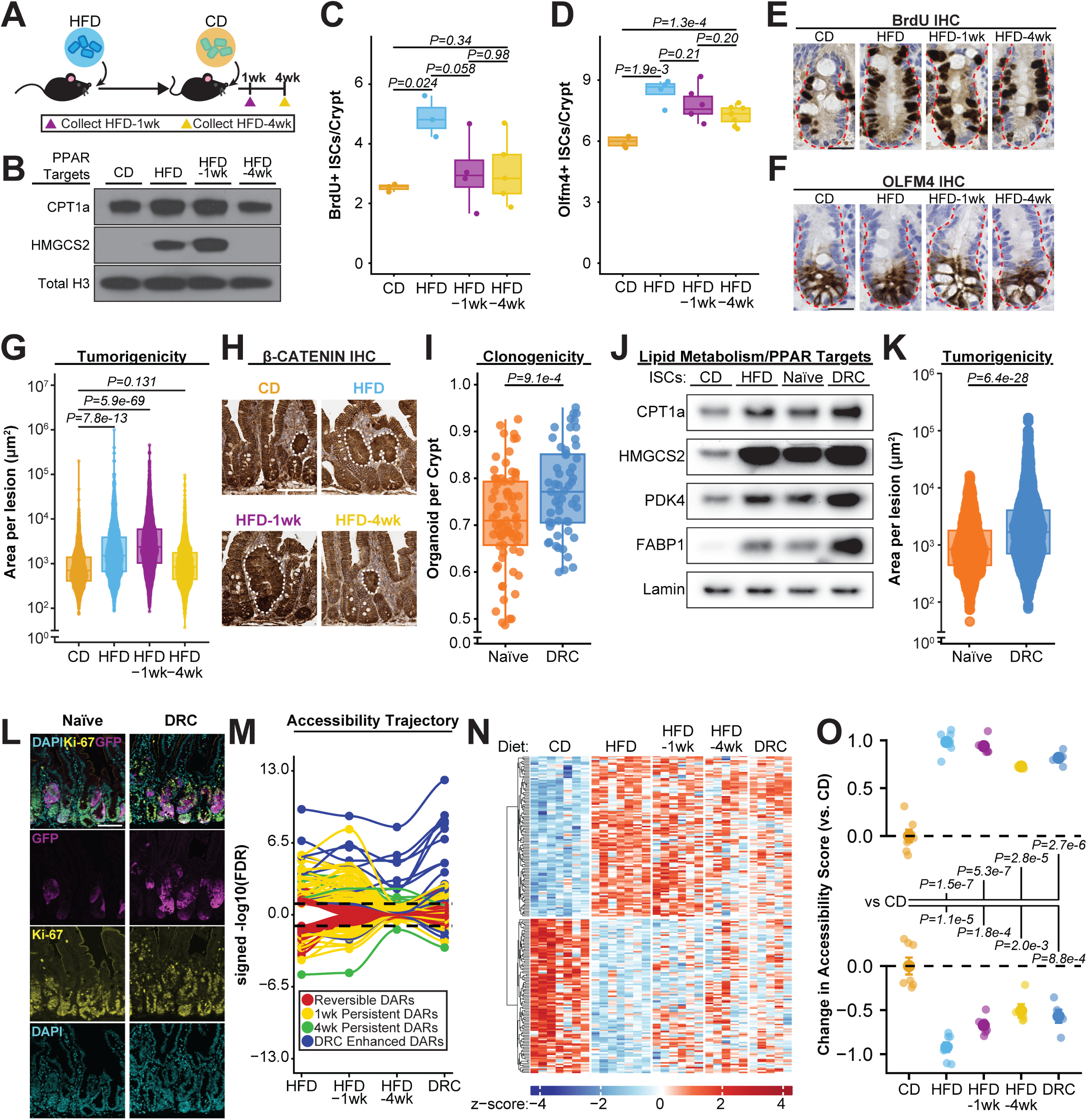
HFD-induced chromatin changes persist following dietary normalization. **(A)** Experimental schematic showing collection of intestinal crypts from mice maintained on HFD for 24 weeks and subsequently returned to CD for 1 week (HFD-1wk) or 4 weeks (HFD-4wk). **(B)** Western blot of flow-sorted EPCAM^+^ intestinal crypt cells showing protein levels of PPAR-target genes CPT1A and HMGCS2 in CD, HFD, HFD-1wk, and HFD-4wk groups. Total H3 serves as loading control (n=2). **(C,D)** Quantification of (**C**) BrdU^+^ and (**D**) OLFM4^+^ ISCs per intestinal crypt across the dietary conditions. Each point represents individual mean of >20 crypts analyzed per mouse (WT ISCs: CD n=3, HFD n=3, HFD-1wk n=4, HFD-4wk n=5). **(E,F)** Representative IHC images showing (**E**) BrdU and (**F**) OLFM4 staining in intestinal crypts in **(C,D)**. Scale bar = 25 µm. **(G)** Quantification of β-catenin^+^ adenomatous lesion area (µm²) measured from CD, HFD, HFD-1wk, and HFD-4wk mice. Each point represents individual lesion area measured from mouse cohorts (Individual *Apc*^KO^ mice: CD n=7, HFD n=7, HFD-1wk n=6, HFD-4wk n=8.) **(H)** Representative images of β-catenin^+^ adenomas from each dietary condition, dotted white line represents adenomatous area within tissue. Scale bar = 100 µm. **(I)** Quantification of clonogenic capacity measured by the number of organoids formed per crypt from naive and DRC ISCs. Each point represents the organoid formation rate per well of plated crypts derived from individual mice. We measured 5 or more wells of plated crypts per mouse for clonogenicity measures. Individual WT mice: Naïve n=12, DRC n=9. **(J)** Western blot of flow-sorted ISCs showing PPARd/a target genes CPT1A, HMGCS2, PDK4, and FABP1 abundance in CD, HFD, naive, and DRC conditions. Lamin serves as the loading control. (n=2) **(K)** Quantification of adenomatous lesion area (µm²) measured from naïve versus DRC intestinal tissues stained with β-catenin. Each point represents the area of an individual lesion from *Apc*^KO^ mice: Naïve n=4, DRC n=6. **(L)** Representative images from (**K**) highlighting adenomas with DAPI-stained nuclei (cyan), *Lgr5*^GFP^ reporter (magenta), and KI-67 proliferation marker (yellow) in naive and DRC intestinal sections. Scale bar = 100 µm. **(M)** Accessibility trajectory analysis showing signed –log_10_FDR values for differentially accessible regions across sequential dietary transitions (HFD, HFD-1wk, HFD-4wk and DRC). **(N)** Heatmap of z-scored accessibility values for genomic regions showing progressive reversion or persistence of accessibility across CD, HFD, HFD-1wk, and HFD-4wk, and DRC conditions. **(O)** Accessibility module scores relative to CD, HFD, HFD-1wk, and HFD-4wk, and DRC groups (two-tailed unpaired *t*-test comparisons to WT CD ISCs; *P* < 0.05).

We tested whether loss of the HFD phenotype is reflected in the underlying chromatin landscape. By augmenting our current CD and HFD data with ATAC-seq of ISCs derived from mice removed from the HFD exposure for 1 and 4 weeks, we find that, generally, the HFD and HFD-1wk samples exhibit similar variance by tighter clustering in the PCA (Figure S5A), while control and HFD-4wk samples cluster closer together, in essence recapitulating observations *in vivo* with recovery from HFD-induced changes. We identified DARs at each time point relative to the CD condition, finding that > 63% of DARs that increase in accessibility due to HFD are retained at HFD-1wk (Figures S5B and S5D), and a retention of 18% at HFD-4wk (Figures S5C and S5E). These remaining peaks are enriched for PPAR and HNF binding sites at both timepoints after diet normalization (Figures S5D and S5E). While these dynamics display a rapid change after removal from the HFD, chromatin accessibility persists well after the regenerative and tumorigenic capacity reverse with accessibility profiles failing to return to the CD state (Figures S5F and S5G). In contrast to HFD-induced open peaks, only ∼17% of DARs that decrease in accessibility due to HFD are still observed at HFD-1wk and less than 1% retained at HFD-4wk (Figure 3M). These data show that the HFD drives increased accessibility that is retained, whereas areas of decreased accessibility rapidly recover to CD levels.

### ISCs remember the HFD

To evaluate whether the persistence of HFD-induced chromatin accessibility confers a heightened capacity for ISCs to respond to renewed HFD exposure, we established a diet re-challenge (DRC) model (Figure S5H). Mice chronically exposed to HFD were removed for 4 weeks to allow for phenotypic reversion then subsequently challenged with the HFD for a 1-week exposure. Mice on a chronic CD with 1-week HFD exposure served as the controls (naïve). We first tested ISC functionality by assessing crypt clonogenicity. Crypts harvested from DRC mice were more clonogenic than the naïve control counterpart (Figure 3I). Molecularly, we see an elevation of lipid metabolism and PPAR targets between the two conditions (Figure 3J). Histologically, OLFM4^+^ cells remains elevated in the DRC intestines relative to naïve controls albeit non-significantly (Figure S5I), and proliferation increases in both populations (Figure S5J) suggesting that the enhanced clonogenicity of DRC crypts is not explained by a difference in cell-cycle rate. We next tested if *Apc* loss, a driver of adenoma formation, would similarly increase adenomatous growth in the DRC mice. Indeed, loss of *Apc* induced after the HFD re-challenge resulted in larger adenomatous β-catenin^+^ lesions and increased Ki-67 proliferation (Figure 3K and 3L) in mice re-exposed to the HFD, altogether indicating that the persistent chromatin state after CD normalization may be the basis for a molecular memory and enhanced response to reintroduction of the HFD.

To assess how the chromatin accessibility facilitates these changes, we evaluated the ISC chromatin in the DRC mice to find an increase in the DARs present relative to CD (no HFD exposure). The DRC-induced changes represent the reestablishment of 26% of the open peaks observed in the original HFD DARs, with 31 new regions differentially accessible (Figure S5K and S5L). Conversely, 7% of closed HFD DARs were observed in the DRC. Moreover, within this HFD subset, the chromatin profile of the DRC samples showed a rebound effect in which regions that had previously begun to close at HFD-1wk or HFD-4wk exhibit amplified accessibility beyond the original levels of chronic HFD-exposed mice (Figure 3M). While DARs from chronic exposure to HFD were relatively similar in number to regions that gained or lost a significant degree of accessibility (Figure 1B), the DRC mice are predominantly biased towards DARs that increase in accessibility. This increase suggests that short-term re-challenge of the HFD drives rapid alterations in chromatin accessibility that favors a more permissive chromatin state rather than promoting chromatin closure. Overall, a brief 1-week HFD re-exposure did not recapitulate a complete HFD-DAR profile, indicating that most changes are likely progressive and accumulate with time (Figures 3N and 3O). This also suggests that the missing DARs in the DRC (Figures S5K and S5L) may not be responsible for the enhanced phenotypic robustness in stem cell self-renewal allowing for other factors to influence the rapid functional response to reintroduction, such as cellular metabolism that can influence ISC function outside of chromatin remodeling, or other priming epigenetic marks not captured in this study. How these later features are programmed in the ISCs remains unclear.

### *Apc* loss masks diet-induced accessibility changes

The HFD leads to increased spontaneous and genetically-induced adenomas, emphasizing the risk in diet-induced obesity (Beyaz et al., 2016; DeClercq et al., 2015; Mana et al., 2021; Niku et al., 2017; Schulz et al., 2014; Yang et al., 2022). Loss of functional *Apc* is the dominant driver mutation in intestinal tumorigenesis (The Cancer Genome Atlas Network, 2012). We leveraged our system to explore the change in chromatin accessibility in the transition of WT to adenomatous ISCs, and the contribution of a HFD on the chromatin accessibility. First, we performed ATAC-seq on *Lgr5*^GFPhi^ ISCs isolated from adenomas (*Apc*^KO^) from mice on a CD or HFD and found that the two *Apc*^KO^ conditions are highly similar across the chromatin landscape (Figure S6A). Furthermore, the two different diets separate the *Apc*^KO^ samples by only a small number of peaks (Figure S6B), thus implying that loss of *Apc* is a normalizing nuclear event and diet has a minimal impact on the chromatin accessibility. However, the comparison between the WT and adenoma transition in *Lgr5*^GFPhi^ cells reveals large changes in accessibility and transcription. When comparing *Apc*^KO^ to the WT counterparts, we find that the *Apc*^KO^ samples are distinct from WT (Figure 4A) with more than 20,000 *Apc*^KO^-driven DARs, exhibiting two orders of magnitude more DARs than observed in WT HFD-exposure (Figures 4B and 4C). To explore the differences between WT and *Apc*^KO^, we directly compared the accessible regions within the CD-fed WT and *Apc*^KO^ ISCs (Figures 4B and 4D) and used published scRNAseq data to further distinguish these two states(Bala et al., 2023) (Figures S6E-G). Accessible regions that increase within promoters in *Apc*^KO^ are associated with diverse ligand-receptor signaling systems, such as the Ras pathway as well as downstream MAPK signaling (Figure 4E). The motif enrichment within the accessible regions is heavily associated with immune and stress response programs (Figure 4F) that include AP-1 (Jun/Fos) and Fra1/2 (Fos-related antigen-1/2), sites which are consistent with Kras-mediated AP1 activation and Fra1-regulated inflammation and immune function (Mzoughi et al., 2025; Klomp et al., 2024; Sen et al., 2019; Shaulian and Karin, 2001; Mechta et al., 1997). Additionally, binding sites for chromatin architecture proteins, CTCF and Boris, are also heavily enriched (Figure 4F), suggesting rewiring of chromatin organization upon loss of a tumor suppressor (Ghirlando and Felsenfeld, 2016; Hnisz et al., 2016; Marshall et al., 2014). These patterns are also present in the HFD-exposed WT to *Apc*^KO^ *Lgr5*^GFPhi^ ISCs accessibility analysis (Figures S6C and S6D). The associated transcriptional increases yield pathways linked with immune response to infections consistent with AP1/Fra activity (Figure S6H). The decreased accessible regions are enriched in programs related to nuclear receptor and RXR activity, whose functions are associated with differentiation such as retinol (RAR), bile (FXR), and arachidonic and fatty acid (PPARs) metabolism (Evans and Mangelsdorf, 2014; Lukonin et al., 2020; Makishima et al., 1999; Modica et al., 2008; Kliewer et al., 1997) (Figure 4G). Transcriptionally, the downregulated programs are enriched for oxidative phosphorylation, consistent with the decrease in accessibility and the association with differentiation (Mzoughi et al., 2025) (Figure S6K). When we explore the potential transcription factor binding motifs within accessible regions, HNF4a, PPAR, and RXR are heavily represented (Figure 4G), suggesting that the *Apc*^KO^ ISCs are less responsive to environmental cues and mitochondrial-mediated energy production, while being more consistent with the early steps in tumor progression (Wester et al., 2021; Bala et al., 2023; Mzoughi et al., 2025).

**Figure 4.**
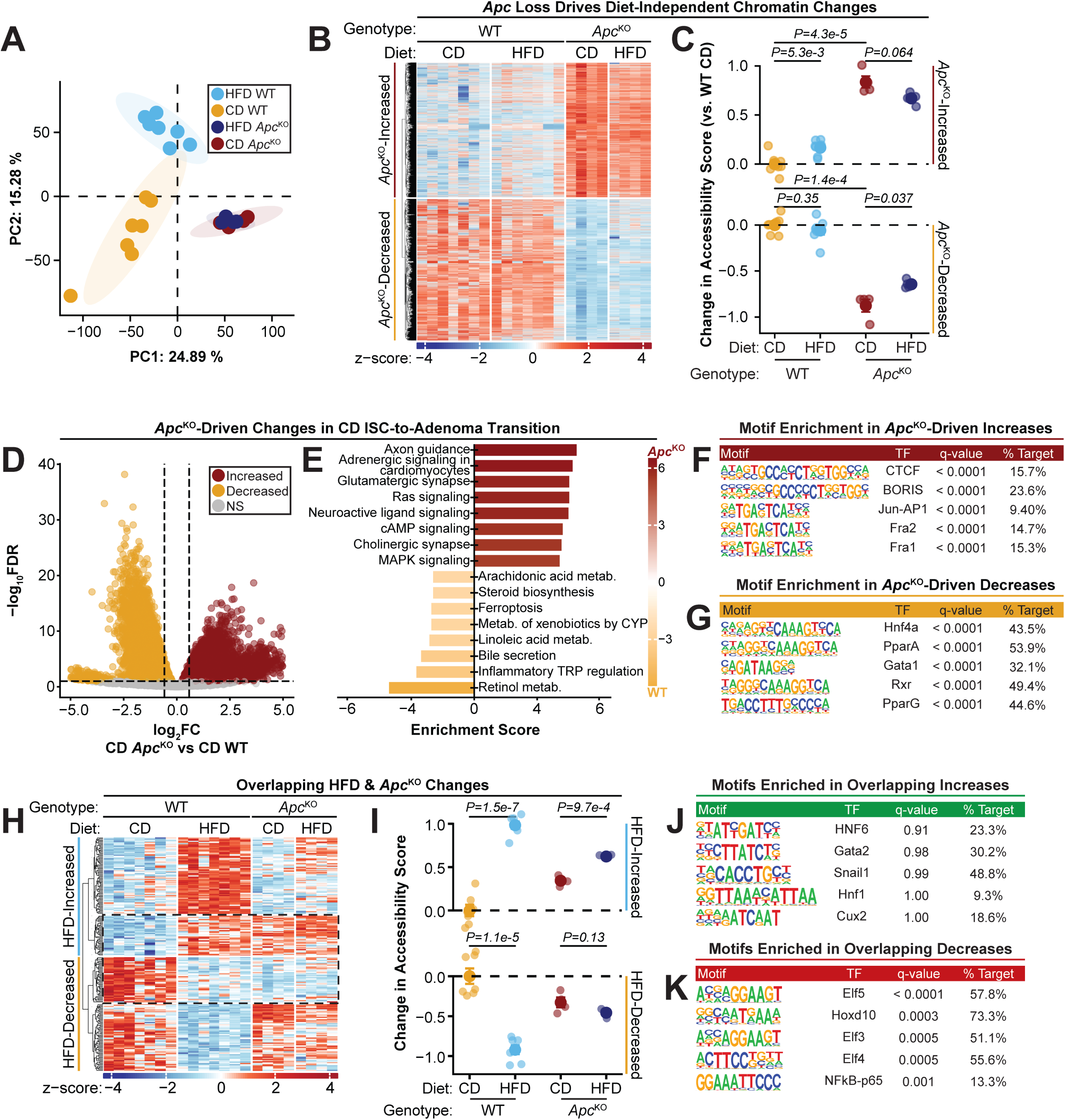
HFD-induced accessibility overlaps with *Apc*^KO^ loci. **(A)** PCA of variance-stabilized chromatin accessibility profiles from *Apc*^WT^ and *Apc*^KO^ ISCs derived from CD and HFD-fed mice. **(B)** Heatmap of z-scored chromatin accessibility values showing regions with increased or decreased accessibility in *Apc*^KO^ relative to *Apc*^WT^ samples across CD and HFD conditions. **(C)** Dot plots displaying accessibility module scores for *Apc*^KO^-increased and *Apc*^KO^-decreased regions in CD and HFD samples (two-tailed unpaired *t*-test). **(D)** Volcano plot showing DARs between *Apc*^WT^ and *Apc*^KO^ ISCs under CD. Each point represents an individual genomic region: orange = increased accessibility, red = decreased accessibility, gray = nonsignificant. Significance threshold: FDR < 0.1; |log_2_fold-change| > log_2_1.5. **(E)** KEGG pathway enrichment bar plot for open (gold) and closed (red) DARs identified in *Apc*^KO^ relative to *Apc*^WT^ ISCs. **(F,G)** Motif-enrichment analysis of **(F)** *Apc*^KO^-increased and **(G)** *Apc*^KO^-decreased regions showing top TF matches, q-values and percent of target loci containing the matching TF motif. **(H)** Heatmap of z-scored accessibility for genomic regions of HFD-induced changes. Dotted lines represent overlapping *Apc*^KO^-induced accessibility changes, separated by increased and decreased groups. **(I)** Dot plots showing accessibility module scores for overlapping increased and decreased regions across CD and HFD samples in *Apc*^WT^ and *Apc*^KO^ genotypes (two-tailed unpaired *t*-test). **(J,K)** Ranked motif-enrichment analyses within overlapping **(J)** increased and **(K)** decreased regions, displaying motif sequence logos, TF matches, q-values, and percent of target loci containing the matching TF motif.

To determine whether the chromatin changes induced by the HFD in WT ISCs overlap with those found in *Apc*^KO^, we compared the WT HFD-induced DARs with the DARs identified in *Apc*^KO^ *Lgr5*^+^ ISCs. The original WT HFD-induced DARs overlap with *Apc*^KO^ DARs with high significance (hypergeometric *P-value* < 1.0^-10^) within a subset of 43 and 45 shared loci in the open and closed directions respectively, representing nearly half the total HFD-induced DARs (Figures 4H-I, S6I, and S6J). This concordance indicates that a substantial portion of the chromatin remodeling promoted by a HFD occurs at regions that are also reconfigured during oncogenic transformation. This subset of the WT-HFD DARs exhibited consistent accessibility patterns between CD and HFD *Apc*^KO^ conditions (Figure 4H and 4I), highlighting that HFD and *Apc*-loss share a common set of regulatory elements that remain stably altered. In contrast to previous HFD-induced regions (Figure 1B and 1D), this subset does not harbor significant enrichment of PPAR binding motifs in the open regions and instead is enriched with fundamental intestinal-specific developmental factors like HNF1 and Snail1, albeit non-significantly(Horvay et al., 2015; Beyes et al., 2019; Yang et al., 2016) (Figure 4J). The closed regions continue to contain significant NFkB factor binding sites, relaying a consistent theme in WT-HFD and *Apc*^KO^ samples (Figure 4K). In summary, HFD induces durable chromatin changes that converge with *Apc*^KO^ remodeling, yet loss of *Apc* fundamentally rewires epigenetic programming, rendering these cells refractory to chromatin changes imposed by diet. Indeed, when we combine all samples to assess global variability across cell types, genotypes, and diet, the *Apc*^KO^ populations appear distinct from their cell-of-origin (Figure S6K), mirroring a similar divergence of TACs, whereas diet and the different genotypes examined in ISCs remain comparatively constrained.

## DISCUSSION

In this study, we examined how ISCs adapt to a high-fat Western diet by profiling changes in chromatin accessibility. Our findings reveal that dietary lipid exposure induces more than transcriptional changes and enhances alterations in the chromatin landscape independent of loci related to stem cell identity. Importantly, HFD-induced chromatin alterations are retained during the transition from ISCs to TAC progenitors, implying that the stem cell epigenome transmits diet-encoded features that may enhance proliferative and regenerative capacity in the progeny. This persistence suggests that the metabolic response can imprint molecular memory within the ISC lineage. Mechanistically, we demonstrate that these chromatin dynamics depend on lipid-sensing nuclear receptors PPARd and PPARa, but not on CPT1A, the canonical enzyme they transcriptionally modulate for mitochondrial long-chain FAO. This distinction underscores that PPAR activation and signaling, rather than downstream metabolic flux, serves as the central axis linking dietary lipids to chromatin remodeling. In this framework, nuclear receptors act as signal-responsive chromatin modulators that integrate nutrient cues. We observe that ISC HFD-induced chromatin accessibility changes persist upon diet normalization despite reversal of overt phenotypes thereby uncoupling the molecular state from the functional recovery and hypothesize that residual modifications may leave the epigenome “primed” for an enhanced response upon re-exposure. The duration and stability of this permissive chromatin state remain unknown but may represent a window during which prior dietary experience shapes future regenerative behavior and disease susceptibility.

The DARs we identified reflect enhanced activation of pre-existing accessible chromatin, rather than the opening of new domains of accessibility (Li et al., 2014; Qin et al., 2018). Even at the most persistent loci, chromatin accessibility was already established, suggesting that a HFD and PPAR nuclear receptor signaling exert limited capacity to pioneer new cis-regulatory elements. Instead, their influence likely occurs through modulation of pre-accessible enhancers. In contrast to PPARs, which act in a ligand-dependent and context-specific manner, PRDM16 and HNF4α/γ serve as obligate regulators of basal lipid metabolism, where their loss results in epithelial degeneration (Chen et al., 2020; Stine et al., 2019). These factors may function upstream or in parallel with PPARs, directing their recruitment to relevant metabolic loci and ensuring that transcriptional responses occur within a pre-defined regulatory landscape. Notably, most HFD-induced DARs and PPARd binding sites were located within distal enhancer and intronic regions, which may not directly confer transcriptional activation. PPARd binds DNA in its unliganded, transcriptionally inactive state, positioning these sites as poised enhancers (Shi et al., 2002). One possibility is that PPARd facilitates higher-order chromatin organization, such as stabilizing enhancer-promoter contact loops or local chromatin boundaries, thereby indirectly modulating gene accessibility rather than directly driving transcription. This mechanism would be consistent with other nuclear receptors. For example, HNF4, an orphan nuclear receptor, has been shown to promote local enhancer-enhancer and enhancer–promoter contacts and maintain chromatin architecture necessary for epithelial function (Chen et al., 2021). Interestingly, steroid hormone receptors such as glucagon, estrogen, and androgen receptors appear to function in similar ways by operating on pre-established chromatin regions (Altıntaş et al., 2024; D’Ippolito et al., 2018; Kocanova et al., 2024). These receptors reside in the cytoplasm and upon activation enter the nucleus. Rather than generating new chromatin loops to remodel chromatin architecture they instead increase the chromatin contact frequency at loops with pre-existing receptor-bound sites, indicating that changes in chromatin contact propensity, as opposed to new loop formation, can carry significant regulatory and biological consequences.

A recurring theme in our analysis is the suppression of immune-modulatory programs under HFD conditions. We find that regions gaining accessibility in HFD ISCs are enriched for PPAR binding motifs, whereas regions losing accessibility show depletion of NFκB binding sites. This inverse pattern suggests that PPAR activation may antagonize NFκB-dependent transcriptional networks, thereby repressing inflammation-associated programs. Such findings align with accumulating evidence that lipid-rich metabolic states suppress immune signaling pathways (Bensinger and Tontonoz, 2008; Garcia et al., 2023; Prendeville and Lynch, 2022). Mechanistically, PPARs and NFκB are known to engage in transrepression, both directly and indirectly. PPARa can inhibit NFκB activity through a physical interaction with the p65 subunit (Delerive et al., 1999), while activated PPARd has been shown to associate with ERK5, enforcing a PPARd/ERK-mediated repression of NFκB signaling (Woo et al., 2006). In addition, PPARg can function as an E3 ubiquitin ligase targeting NFκB for degradation (Hou et al., 2012), providing a direct molecular mechanism for regulation between these pathways. These modes of crosstalk suggest that direct PPAR–NFκB interactions could similarly occur within ISCs, contributing to the observed immune suppression under HFD. Alternatively, PPARs may influence inflammatory modulators indirectly through metabolic or microbial intermediates. For example, *Beyaz et al.* reported that in HFD ISCs, immune modulation arose from HFD-induced microbiome changes that altered epithelial-immune interactions. In this view, the NFκB-associated chromatin regions that become inaccessible in our dataset may reflect secondary consequences of microbiome-driven signaling alterations rather than direct nuclear receptor activity. Together, these observations highlight a complex interaction in which PPAR activation and NFκB suppression jointly refine the immune-suppressed chromatin state of ISCs under metabolic stress. Whether this arises from direct transcriptional cross-regulation or indirect modulation through the epithelial-microbial axis remains an open question warranting further investigation.

An important question we sought to address was whether ISCs exposed to an over-nutrient state retain features after diet normalization which confers heightened responsiveness upon secondary dietary challenge. This concept of diet-induced stem cell memory posits that metabolic exposures may influence long-term tissue behavior, providing a potential epigenetic basis for the persistent effects of diet on regeneration and disease susceptibility (Hinte et al., 2024; Novak et al., 2021). Evidence of stem cell memory comes from other systems. Skin stem cells exposed to injury exhibit a form of epithelial trained immunity, wherein repair-associated genes respond more robustly to subsequent inflammatory assaults (Gonzales et al., 2021; Larsen et al., 2021; Naik et al., 2017). Similarly, hematopoietic stem cells exposed to β-glucan or lipopolysaccharide become primed for secondary stimulation, showing rapid reactivation of metabolic and proliferative gene programs upon re-exposure (De Laval et al., 2020; Mitroulis et al., 2018). In contrast to these injury or infection-driven contexts, where regeneration and recovery may necessitate chromatin remodeling, the dietary perturbation we examined elicits a more subtle, but persistent, chromatin effect. We observe sustained accessibility of HFD-induced peaks following diet withdrawal accompanied with moderate recovery after HFD-reintroduction. ISCs re-exposed to the HFD exhibited a more robust functional response compared to naïve ISCs, suggesting that the chromatin landscape retains a latent imprint of prior metabolic exposure. Although we did not directly map these features, our findings imply that dietary experience leaves behind an epigenetic “memory trace”, a poised state that enhances responsiveness upon re-challenge.

Given the established link between a Western diet and increased risk for colorectal cancer, we hypothesized that HFD exposure might elicit chromatin accessibility patterns similar to those preceding oncogenic transformation. Indeed, a large fraction of the HFD-induced DARs are significantly overlapping in *Apc*^KO^ loci and few of these loci were observed in the fasted state. Unexpectedly, however, loss of *Apc* profoundly rewires the chromatin accessibility and largely ignores HFD exposure after transformation. The magnitude of chromatin remodeling accompanying the transition from WT to *Apc*^KO^ ISCs far exceeded changes induced by dietary perturbation, underscoring that oncogenic transformation is a dominant nuclear event for chromatin reorganization relative to metabolic input. Similar to the HFD-*Cpt1a*^iKO^ accessibility, cytoplasmic metabolic shifts are buffered in the nucleus. Interestingly, among the loci most affected by *Apc*-loss were those related to RXR, the obligate heterodimeric partner for PPARs and numerous other nuclear receptors (Evans and Mangelsdorf, 2014), which exhibited significant decreases in chromatin accessibility. This finding highlights a potential mechanism by which *Apc*-loss may suppress RXR-mediated transcriptional programs, thereby blocking differentiation, reducing oxidative metabolism, and preserving stem-like identity within the epithelium (Bala et al., 2023; Mzoughi et al., 2025; Wester et al., 2021). Intriguingly, while HFD exposure enhances PPAR:RXR signaling activity, *Apc*-loss appears to exert the opposite effect, suggesting a mechanistic tension between metabolic activation and oncogenic repression in shared nuclear receptor networks. This apparent divergence raises the question of how *Apc*^KO^ adenomas use lipids in this context and how the cellular metabolism rewires to accommodate *Apc*-loss and leverage a HFD for growth (Beyaz et al., 2016; Mana et al., 2021). Ultimately, *Apc*-loss reveals that environmental inputs such as an HFD, have minimal impact on stem cell chromatin states with cytoplasmic metabolism buffering the chromatin, and that tumor suppressor genetic lesions exert a dominant and enduring control over epigenetic identity, highlighting a complex interplay between diet, metabolism, and oncogenic signaling in the early stages of colorectal cancer.

## METHODS

### Experimental model and subject details

Mice were under the husbandry care administered by Arizona State University’s Department of Animal Care and Technologies. All procedures were conducted in accordance with the American Association for Accreditation of Laboratory Animal Care and approved by ASU’s Institutional Animal Care and Use Committee. The following alleles were obtained from the Jackson Laboratory or gifted: *Ppard*^fl/fl^ (B6.129S4-Ppard^tm1Rev^/J, stock number 005897(Barak et al., 2002)); *Ppara*^−/−^ (129S4/SvJae-Ppara^tm1Gonz^/J, stock number 003580(Lee et al., 1995)); *Lgr5*^eGFP-IRES-CreERT2^ (B6.129P2-*Lgr5^tm1(cre/ERT2)Cle^*/J(Barker et al., 2007)); *Cpt1a*^fl/fl^ (Gift from Peter Carmeliet(Schoors et al., 2015)); *Villin*^CreER^ (B6.Cg-Tg(Vil1-Cre/ERT2)23Syr/J(el Marjou et al., 2004)); *Apc*^fl/fl(exon^ ^14)^ (*Apc*^fl/fl^) has been previously described(Colnot et al., 2004). The following strains were bred in-house: 1) *Villin*^CreER^; *Lgr5*^eGFP-IRES-CreERT2^, 2) *Ppard*^fl/fl^; *Ppara*^−/−^; *Villin*^CreER^; *Lgr5*^eGFP-IRES-CreERT2^, 3) *Cpt1a*^fl/fl^; *Villin*^CreER^; *Lgr5*^eGFP-IRES-CreERT2^, 4) *Apc*^fl/fl^; *Lgr5*^eGFP-IRES-CreERT2^. Mice were sex and age-matched per experiment and housed under a 12/12 day/night light cycle. Floxed alleles were excised following intraperitoneal (IP) injection of tamoxifen, administered at 25-100 mg/kg. Mice harboring *Ppar* or *Cpt1a* alleles were administered tamoxifen 3X at 50 mg/kg prior to establishment on diet. BrdU (Sigma Aldrich, 19-160) was prepared at 10 mg/ml in PBS and IP injected at 100 mg/kg 4 hours prior to harvesting tissue. High-fat diet (Research Diets D12492) and control diet (matching sucrose, Research Diets D12450J) were provided to mice *ad libitum* at the age of 8-12 weeks for 6 to 8 months.

### Crypt isolation and culture

The small intestines were removed, flushed with PBS^−/−^ + 10 mM EDTA, opened, gently wiped to remove mucus layer, and cut into ∼3 cm sections. Intestine pieces were rinsed 3X and incubated on ice for 30 minutes. Crypts were mechanically separated from the connective tissue by shaking and filtered through 70 μm mesh into a 50 mL conical tube to remove villi. Isolated crypts were counted and embedded in 4:6 media:Matrigel (Corning 356231 growth factor reduced) mixture at 5-10 crypts per μl density and cultured in a medium modified from Sato et al. (2009)(Sato et al., 2009) using the following: Advanced DMEM (Gibco, 12491015), Glutamax (Gibco, 35050-061), Penicillin:Streptomycin (VWR, 97063-708), 40 ng/ml EGF (Peprotech, 315-09), conditioned media approximate to 200 ng/ml Noggin and 250 ng/ml R-Spondin, 1 μM *N-*acetyl-L-cysteine (Sigma Aldrich, A9165), 1XB27 (Life Technologies, 17504044), 1 μM Chir99021 (LC Laboratories, C-6556), and 10 μM Y-27632 dihydrochloride monohydrate (Sigma Aldrich, Y0503). Cultures were maintained at 37°C and 5% CO_2_.

### Isolation of ISCs/TACs

Crypt suspensions were pelleted (300 g, 5 minutes, 4°C) and resuspended in TrypLE (Gibco, 12604039) and dissociated by heat shock at 32°C for 90 seconds. Filtered single cells were stained with the following antibody cocktail for flow cytometry analysis: CD45-PE (BioLegend, 103106, 1:500, 10 minutes, pan-immune marker), EpCAM-APC (Invitrogen, 17-5791-82, 1:300, 10 minutes, pan-epithelial selection) and DAPI (BioLegend, 422801, 1μg, live/dead marker). Additional gating separated out ISC (*Lgr5*^GFPhi^) and TAC (*Lgr5*^GFPlow^) populations.

### Western Blotting

Crypt epithelial cells (EPCAM-APC⁺) or ISCs (*Lgr5*^GFPhi^) were sorted directly into Laemmli sample buffer (Alfa Aesar, J61337). Per experiment, equal number of cells per sample were loaded onto 4%–12% gradient gels (Invitrogen, NP0335BOX) and probed with the following primary antibodies: Anti-FABP1 (CST, Rabbit mAb, 1:1,000, Cat#13368); Anti-HMGCS2 (Abcam, Rabbit mAb, 1:500, Cat#ab137043); Anti-CPT1A (Abcam, Mouse mAb, 1:500, Cat#ab128568); Anti-PDK4 (Abcam, Mouse mAb, 1:1,000, Cat#ab214938); Anti-Lamin B1 (Abcam, Rabbit mAb, 1:2,000, Cat#ab133741); Anti-Total H3 (CST, Rabbit mAb, 1:10,000, Cat#4449S). Secondary antibodies were used at a 1:3,000 dilution: HRP-linked Anti-Rabbit IgG (CST, Cat#7074S); HRP-linked Anti-Mouse IgG (CST, Cat#7076S).

### Immunostaining

Small intestines were washed with cold 1X PBS, flushed with 10% buffered formalin, fixed at room temperature for 24 hrs, then transferred to 70% ethanol for embedding. FFPE tissue were sectioned (5μm). Antigen retrieval was performed using 1X Borg Decloaker (BioCare Medical, Cat#BD1000G1). The following antibodies were used for Immunofluorescence and Immunohistochemistry (IF/IHC): anti-Ki67 (Invitrogen, Recombinant Rat mAb SolA15, 1:500, Cat#740008T); anti-GFP (Abcam, Goat pAb, 1:500 Cat#ab6673), anti-BrdU (Abcam, Rat mAb, Cat#ab6326); anti-OLFM4 (CST, Rabbit mAb, Cat#39141S). IF secondary antibodies: AF488 (Invitrogen, Alexa Fluor™ 488, 1:1,000, Cat# A-21208) or AF647 (Invitrogen, Alexa Fluor™ 647, 1:1,000, Cat# A-21447) for GFP at a 1:1,000 dilution. DAPI at 1:1,000. Images were captured using a Keyence BZ-X800. The following antibodies were used for immunohistochemistry (IHC): **ATAC-seq Library Prep, Alignment, Trimming, and Peak Calling**

ATAC-seq libraries were prepared on flow-sorted cells (35,000-40,000 per reaction) obtained from individual mice following an adapted protocol from (Buenrostro et al. 2013), generating one library per mouse (biological replicates = number of mice per condition, no pooling of mice).

Sequencing was carried out on an Illumina NextSeq2000 platform (MIT BioMicro Center) using a 40 + 40 base pair-end run with dual indexing (6 + 6 nucleotide indexes). Raw sequencing data were quality assessed with FastQC. Adapter sequences were removed using NMerge, and trimmed sequences were aligned to the mouse reference genome (mm10) using Bowtie2 (v2.4.2) with the --very-sensitive option. Aligned reads were processed using samtools (v1.12) to remove mitochondrial reads and convert file formats from FASTQ to BAM. Duplicate reads were marked and removed using Picard tools (v2.9.0). MACS3 (v3.0.1) was used for peak calling on individual samples, and bedtools2 (v2.31.1) facilitated peak overlap analyses and merging across samples. A minimum of 10 million reads per sample was targeted.

### Selection of Enhancer Regions and Transcription Factor Binding Motifs

Enhancers were determined using differentially accessible genomic regions (DARs), determined by a log2Fold-Change cutoff equivalent to 1.5x accessibility and a False Discovery Rate (FDR) < 0.1. DARs were used for input into HOMER. Motif enrichment was determined using *findMotifsGenome.pl* using the mm10 alignment and the argument for *-size given*. Results from the *knownResults* output were used for downstream interpretation and output was used for determining enriched candidate transcription factor binding sites (TFBSs) across positively and negatively enriched DARs.

### ATAC-seq Differential Accessibility Analysis

All statistical analyses used R (v4.4.0) with Bioconductor packages. Raw read counts per consensus peak read from output consensus bed file counts were produced from MACS3 and bedtools2. To remove batch effects, counts were corrected with the *ComBat_seq* function from the *sva* package, modeling diet, sex, genotype, and cell type as biological covariates. Differential accessibility was assessed using *DESeq2*. We constructed a DESeqDataSet with design ∼ diet + sex + genotype + sequencing_batch + cell_type and applied a Wald test. To facilitate pairwise comparisons, we combined grouping factor group = paste0(diet, cell_type, genotype), refit the model with ∼ group, and reran the Wald test. For each contrast (e.g., HFD_ISC_WT vs. Control_ISC_WT), we extracted results and determined significantly differential peaks were defined as those with FDR < 0.1 and |log2FC| > log2(1.5). These criteria guided all downstream subset analyses (e.g., “open” vs. “closed” peaks). Multivariate analyses included principal component analysis (PCA) on the top 10,000 most variable peaks in either direction using *prcomp*() on variance-stabilized counts. Peak annotation was performed with *ChIPseeker’s annotatePeak*(), defining promoters as ±2 kb around the transcriptional start site (TSS) and assigning genomic features via the UCSC mm10 *TxDb* and *Org.Mm.eg.db* databases. Additional data and package information for downstream analyses in R can be found in our public lab GitHub.

### ATAC-seq Module Score

Module scores were derived to quantify the aggregate accessibility of predefined sets of HFD-responsive peaks. Peaks that significantly (FDR < 0.1 and |log2FC| > log2(1.5)) increased or decreased in accessibility in HFD ISCs (relative to a control condition [i.e. CD ISCs]) were defined as the HFD-increased and HFD-decreased modules, respectively. For each module, normalized ATAC-seq counts (DESeq2 normalized counts) were extracted for all peaks belonging to that module. For each sample, a module score was calculated as the mean accessibility across all peaks in the module. To allow comparison across sample conditions, scores were centered by subtracting the corresponding mean module score of CD ISC samples. This centering provides relative scores in which positive mean values indicate greater accessibility than the mean CD ISCs and negative values indicate lower accessibility than the mean of CD ISCs. Centered module scores were used for downstream visualization and statistical analyses. Heatmaps were generated from the normalized accessibility count matrix prior to centering while all quantitative comparisons between conditions were performed using the centered module scores.

## RESOURCE AVAILABILITY

### Lead contact

Further information and request for resources and reagents should be directed to the lead contact Miyeko D Mana

### Materials availability

This study did not generate new unique reagents.

### Data and code availability

The GEO accession will be provided upon publication.

All further requests should be submitted to the lead contact.

## ACKNOWLEDGEMENTS

We thank Lindsay M LaFave for initial assistance with the ATAC-seq protocol; Ömer H Yilmaz for the use of the initial ATAC-seq data set, *Apc*^KO^-induced reversal data, and the generous gift of mouse strains; Noah Snyder-Mackler and Christopher Plaisier for unlimited guidance on analysis. Cell sorting was assisted by Adam Kindelin and the ASU Biosciences Flow Cytometry Core using BD FACSymphony S6, acquired by the NIH SIG award 1-S10-OD032287-01. ASU Research Computing provided HPC and storage support(Jennewein et al., 2023). BBB is supported by NIBIB-R21EB034970, NIMH-DP2MH136493. MDM is supported by NCI-K22CA241083, RSG-22-095-01-CCB, NCI-R01CA301086, NIDDK-R01DK143944.

## AUTHOR CONTRIBUTIONS

D.R.S.: Conceptualization, Methodology, Software, Validation, Formal analysis, Investigation, Writing—original draft. Y.B.M.: Methodology, Investigation. T.H.M.: Investigation. G.C.: Methodology, Investigation. S.S.: Investigation. E.U.: Investigation. F.F.: Investigation. G.L.: Investigation. K.L.: Writing—review & editing. K.M.: Investigation. B.B.B.: Conceptualization, Methodology, Investigation, Supervision, Writing—review & editing. M.D.M.: Conceptualization, Methodology, Validation, Investigation, Supervision, Writing—original draft, Funding acquisition.

