## Supplemental Data for "Diet-induced chromatin states influence intestinal stem cell memory"

#### SUPPLEMENTAL FIGURE LEGENDS

**Supplementary Figure 1. ISC chromatin accessibility under a HFD.**

**(A)** Representative Western blots of histone modifications from FACS-sorted EPCAM$^{+}$ (crypt epithelia) and GFP$^{hi}$ (ISCs) populations showing H3K27ac, H3K9ac, H3K27me3, and H3K4me levels.

**(B)** Quantification of relative densitometry for ISC-GFP$^{hi}$ histone modifications normalized to total H3.

**(C)** Experimental schematic outlining diet intervention, crypt isolation, single-cell dissociation, and FACS sorting of GFP$^{hi}$ ISCs and GFP$^{low}$ TACs from *Lgr5*$^{eGFP\text{-}IRES\text{-}CreER}$;*Villin*$^{CreERT2}$ mice on CD or HFD.

**(D)** Average signal profiles and heatmaps for the 10,000 most open and 10,000 most closed genomic regions centered around the TSS for CD and HFD ISCs.

**(E,F)** KEGG pathway **(E)** and GO biological process **(F)** enrichment bar plots for open and closed genomic regions identified in CD versus HFD ISCs.

**(G)** Scatter plots comparing ATAC-seq and RNA-seq log$_2$FC values for genes grouped by TSS proximity (0–1 kb, 1–2 kb, 2–3 kb, 3–4 kb, and 4–5 kb away from the TSS).

**(H)** Genome-browser tracks displaying accessibility around Stem Cell genes (*Olfm4*, *Ascl2, Axin2)*, Lipid Metabolism genes (*Fabp1*, *Acaa1b*, *Cpt2)*, and Immunomodulatory gene loci (*Gbp10, Nlrp9b, Mir29b-2)*. Accessibility of individual loci is quantified by sample condition shown below tracks with *P*-values from two-tailed unpaired *t*-tests with multiple-comparison correction performed using the Benjamini–Hochberg FDR method.

**Supplementary Figure 2. Comparison of HFD and fasting accessibility changes.**

**(A)** Heatmap of z-scored accessibility values across HFD-increased and HFD-decreased DARs in ISCs with age-matched CD, HFD, and 24-hour Fasted conditions (WT ISCs: CD n=7, HFD n=7, Fasted n=4).

**(B)** Dot plots displaying accessibility module scores for ISC HFD-increased and HFD-decreased DARs relative to CD with *P*-values from two-tailed unpaired *t*-tests. Each point represents one mouse-derived ATAC-seq sample.

**(C)** PCA plot of variance-stabilized chromatin accessibility values for ISCs under CD, HFD, and 24-hour Fasted conditions. Each point represents one mouse-derived ATAC-seq sample. Point shape distinguishes male/female.

**Supplementary Figure 3. Comparison of differentiation and diet-induced chromatin accessibility changes.**

**(A)** Heatmap of z-scored chromatin accessibility values (counts) for regions differentially accessible between CD ISCs and CD TACs, grouped as increased or decreased with ISC-to-TAC differentiation (WT ISCs: CD n=7, HFD n=7; WT TACs: CD n=4, HFD n=4).

**(B)** KEGG pathway enrichment bar plot for open (green) and closed (red) regions between ISCs and TACs, irrespective of diet.

**(C)** Scatter plots showing correlations between ATAC-seq and RNA-seq log$_2$FC values for genes with promoter accessibility differences (0–1 kb and 1–2 kb from the TSS) in ISCs versus TACs, irrespective of diet.

**(D)** Volcano plot of DARs comparing CD TACs to HFD TACs. Each point represents a genomic region: green points = increased accessibility in HFD, orange points = decreased accessibility, and gray = nonsignificant. Significance threshold: FDR < 0.1; $|\log_2 FC| > \log_2 1.5$.

**(E)** KEGG pathway enrichment bar plot for open (green) and closed (orange) regions in HFD versus CD TACs.

**(F)** Scatter plots of ATAC-seq and RNA-seq $\log_2$FC values for genes near promoter regions (0–1 kb and 1–2 kb from the TSS) in HFD versus CD TACs.

**(G)** Heatmap of z-scored accessibility values for HFD-increased and HFD-decreased DARs identified in ISCs, visualized across ISCs and TACs under CD and HFD conditions (WT ISCs: CD n=7, HFD n=7; WT TACs: CD n=4, HFD n=4).

**(H)** Dot plots displaying accessibility module scores for HFD-increased and HFD-decreased regions across ISCs and TACs. *P*-values from two-tailed unpaired *t*-tests.

**(I,J)** Motif-enrichment analysis for HFD-increased regions **(G)** and HFD-decreased **(H)** regions in TACs displaying the ranked enriched TF motifs, corresponding TF matches, q-values, and percent of target loci containing the matching TF motif.

**Supplementary Figure 4. PPAR signaling contributes to HFD-associated chromatin accessibility and transcriptional activity in ISCs.**

**(A)** Genome-browser tracks displaying chromatin accessibility at representative loci grouped by function. Stem cell-associated genes (*Lgr5*, *Olfm4*, *Ascl2*) and immunomodulatory genes (*Wif1*, *Ltbp1*, *Il23r*) are shown for CD and HFD conditions in WT and *Ppar-d/a*$^{iKO}$ samples. Distances to TSS or intronic positions are indicated above each locus.

**(B)** Pairwise comparison plot showing the relationship of $-\log_{10}$FDR values for differential accessibility between HFD and CD in WT versus *Ppar-d/a*$^{iKO}$ samples.

**(C)** Heatmap of z-scored chromatin accessibility values for HFD-increased and HFD-decreased regions identified in WT mice, shown across CD and HFD conditions in WT, *Cpt1a*$^{iKO}$, and *Ppar-d/a*$^{iKO}$ genotypes (WT ISCs: CD n=7, HFD n=7; *Cpt1a*$^{iKO}$ ISCs: CD n=4, HFD n=4; *Ppar-d/a*$^{iKO}$ ISCs: CD n=6, HFD n=6).

**(D)** Dot plots displaying module scores corresponding to HFD-increased and HFD-decreased regions across the sample types shown in **(C)** (two-tailed unpaired *t*-test against CD ISCs; WT ISCs: CD n=7, HFD n=7; *Cpt1a*$^{iKO}$ ISCs: CD n=4, HFD n=4; *Ppar-d/a*$^{iKO}$ ISCs: CD n=6, HFD n=6).

**Supplementary Figure 5. Chromatin accessibility and functional recovery following HFD withdrawal and dietary re-challenge.**

**(A)** PCA plot of variance-stabilized accessibility profiles for ISCs from CD, HFD, HFD-1wk, and HFD-4wk samples (WT ISCs: CD n=7, HFD n=7, HFD-1wk n=6, HFD-4wk n=5).

**(B,C)** Volcano plots of DARs comparing **(B)** HFD-1wk and **(C)** HFD-4wk versus CD. Each point represents a genomic region: purple = regions with increased accessibility at HFD-1wk, yellow = regions with increased accessibility at HFD-4wk, orange = regions with decreased accessibility, and gray = nonsignificant. Significance threshold: FDR < 0.1; $|\log_2 \text{fold-change}| > \log_2 1.5$.

**(D,E)** Motif-enrichment analysis of regions showing HFD-driven increases after **(D)** 1-week or **(E)** 4-weeks off of HFD. Analysis displays the significance ranked enriched TF motifs, TF matches, and percent of target loci containing the matching TF motif.

**(F,G)** Average signal profiles and heatmaps showing **(F)** increased and **(G)** decreased accessibility centered on the HFD-induced DARs.
**(H)** Experimental schematic outlining the naïve and DRC diet schemes.
**(I,J)** Quantification of **(I)** BrdU$^{+}$ and **(J)** OLFM4$^{+}$ ISCs per crypt across CD, HFD, naïve, and DRC conditions.Each point represents quantified averages of > 20 crypts per one mouse. (WT: CD n=4, HFD n=4, Naïve n=5, DRC n=6).
**(K)** Volcano plot of DARs comparing DRC versus CD groups. Each point represents an individual genomic region: blue = increased accessibility, orange = decreased accessibility, and gray = nonsignificant. Significance threshold: FDR < 0.1; |$\log_2$ fold-change| > $\log_2$ 1.5.
**(L)** Stacked bar plot showing the frequency of DARs that were recovered (dark orange/blue) or not recovered (light orange/blue) following DRC compared with CD.

**Supplementary Figure 6. Integrated chromatin and transcriptional analyses reveal pronounced effects in *Apc*$^{KO}$ with convergence of loci between adenomatous and HFD-derived ISCs.**
**(A)** PCA plot of variance-stabilized chromatin accessibility values from ISCs of *Apc*$^{KO}$ mice under CD and HFD conditions. Each point represents one mouse-derived ATAC-seq sample (*Apc*$^{KO}$ ISCs: CD n=4, HFD n=4).
**(B)** Volcano plot of DARs comparing HFD versus CD *Apc*$^{KO}$ ISCs. Each point represents a genomic region: dark blue = increased accessibility, maroon = decreased accessibility, gray = nonsignificant.
**(C)** Volcano plot of DARs comparing HFD WT versus HFD *Apc*$^{KO}$ ISCs. Each point represents a genomic region: dark blue = increased accessibility in HFD *Apc*$^{KO}$, cyan = decreased accessibility, gray = nonsignificant. Significance threshold: FDR < 0.1; |$\log_2$fold-change| > $\log_2$1.5) **(B,C)**.
**(D)** KEGG pathway enrichment bar plot for open (purple) and closed (cyan) DARs identified in HFD *Apc*$^{KO}$ relative to HFD WT ISCs.
**(E,G)** UMAP plots of the publicly available single-cell RNA-seq dataset GSE224679 showing (**E**) global clustering of *Apc*$^{WT}$ and *Apc*$^{KO}$ ISCs, (**F**) ISC module scores, and (**G**) ISC subsets grouped by genotype.
**(H)** Molecular Signature Database (MSigDB) Hallmark pathway enrichment plots for *Apc*$^{KO}$ up-regulated and *Apc*$^{KO}$ down-regulated gene set. Statistical significance was evaluated using a signed −log10(p-value), where the sign reflects the direction of the corresponding log2 fold change.
**(I,J)** Venn diagrams showing overlap of HFD and *Apc*$^{KO}$-driven accessibility changes for **(I)** increased and **(J)** decreased regions, including number of overlapping peaks and associated significance values.
**(K)** PCA plot of consensus peak counts across all conditions and genotypes, colored by experimental group and shaped by sex.

#### SUPPLEMENTAL METHODS

##### RNA-seq integration

RNA-seq data counts were obtained from GSE151047[10]. We use *biomaRt* package for gene annotation and DESeq2 for downstream normalization and differential expression. DE of genes in each contrast was defined as those with FDR < 0.1 and |log2FC| > log2(1.5). Integration with ATAC-seq data was determined based on expression of genes and presence of accessible genomic region ±2 kb around the TSS. Correlation of accessibility (derived from ATAC-seq analyses) and expression (derived from Bulk RNA-seq analyses) was determined using DE genes and comparing relative log2FC changes at accessible genomic regions in 1kb increments from the TSS.

##### Single-cell RNA-sequencing data processing and integration

Previously published GEO scRNA-seq datasets (GSE164832, GSE151047, GSE224679) were downloaded via GEOquery and analyzed in R (v4.4.0) with Seurat (v5.0). For GSE164832, processed count and metadata tables were extracted directly from the supplementary files. For GSE151047 and GSE224679, count matrices were retrieved from 10x Genomics-formatted files (matrix.mtx.gz, features.tsv.gz, and barcodes.tsv.gz) and parsed with Matrix and data table.

Each dataset was independently converted into a Seurat object using CreateSeuratObject(), with low-quality cells filtered out based on gene detection and mitochondrial content thresholds (nFeature_RNA ≥ 200, percent.mt ≤ 20%). For the GSE224679 dataset, which includes antibody-derived hashtags (HTOs), demultiplexing was performed with HTODemux() (positive.quantile = 0.99), retaining only singlet cells for downstream analyses. Each dataset was normalized using SCTransform (vst.flavor = "v2", method = "glmGamPoi") with mitochondrial read percentage regressed out. Dimensionality reduction was performed using PCA followed by Uniform Manifold Approximation and Projection (UMAP) on the top 30 principal components. Neighbor graph construction and unsupervised clustering were performed with FindNeighbors() and FindClusters() (resolution = 0.2).

### Figure S1

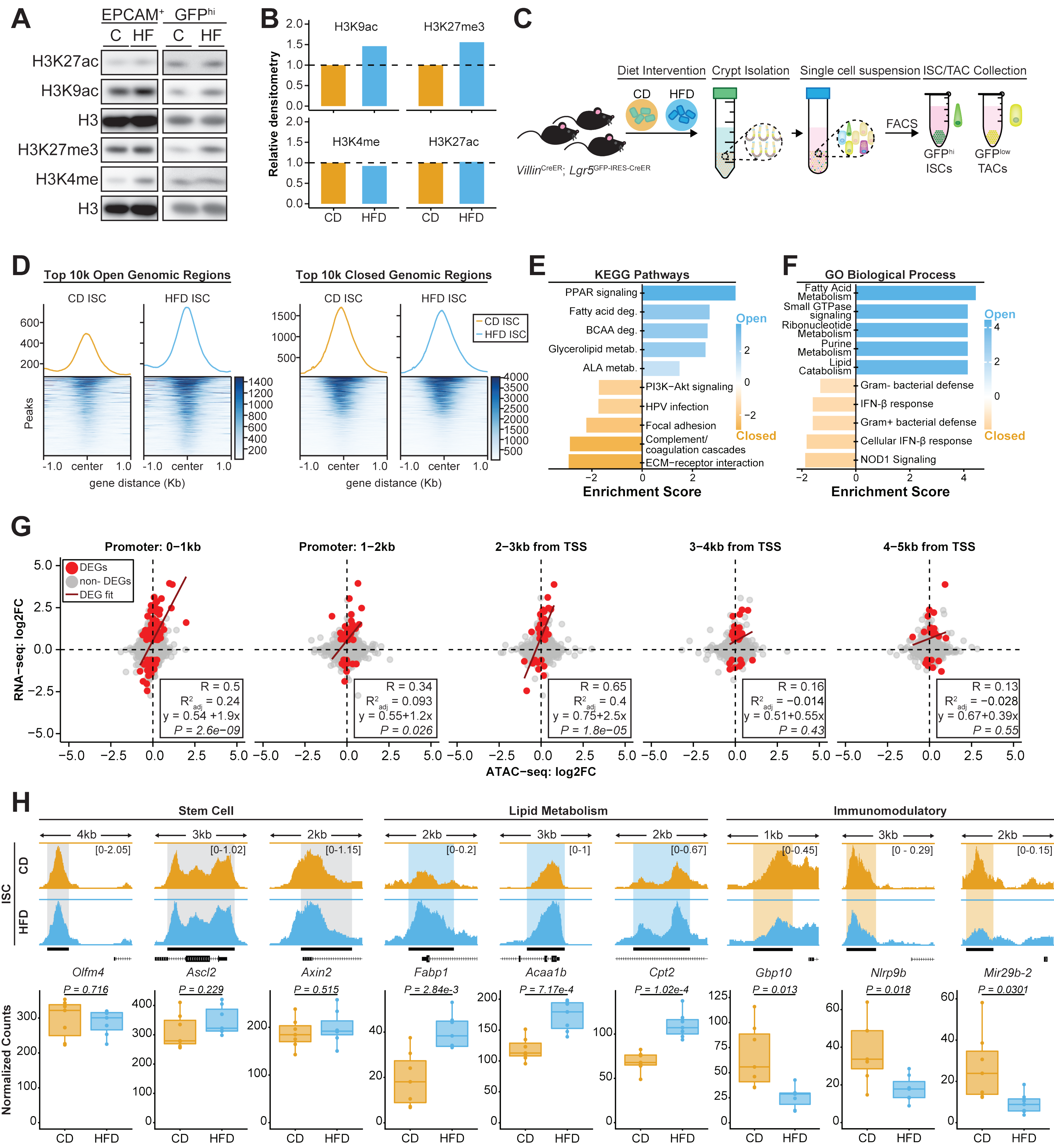

A
EPCAM+ GFPhi
C HF C HF
H3K27ac
H3K9ac
H3
H3K27me3
H3K4me
H3
B
Relative densitometry
H3K9ac
H3K27me3
H3K4me
H3K27ac
CD HFD CD HFD
C
Diet Intervention
Crypt Isolation
Single cell suspension
ISC/TAC Collection
CD HFD
FACS
GFPhi ISCs
GFPlow TACs
VillinCreER; Lgr5GFP-IRES-CreER
D
Top 10k Open Genomic Regions
Top 10k Closed Genomic Regions
CD ISC
HFD ISC
Peaks
gene distance (Kb)
E
KEGG Pathways
PPAR signaling
Fatty acid deg.
BCAA deg.
Glycerolipid metab.
ALA metab.
PI3K−Akt signaling
HPV infection
Focal adhesion
Complement/ coagulation cascades
ECM−receptor interaction
Open
Closed
Enrichment Score
F
GO Biological Process
Fatty Acid Metabolism
Small GTPase signaling
Ribonucleotide Metabolism
Purine Metabolism
Lipid Catabolism
Gram- bacterial defense
IFN-β response
Gram+ bacterial defense
Cellular IFN-β response
NOD1 Signaling
Enrichment Score
G
Promoter: 0−1kb
Promoter: 1−2kb
2−3kb from TSS
3−4kb from TSS
4−5kb from TSS
DEGs
non- DEGs
DEG fit
RNA−seq: log2FC
ATAC−seq: log2FC
R = 0.5 R2adj = 0.24 y = 0.54 +1.9x P = 2.6e-09
R = 0.34 R2adj = 0.093 y = 0.55+1.2x P = 0.026
R = 0.65 R2adj = 0.4 y = 0.75+2.5x P = 1.8e−05
R = 0.16 R2adj = −0.014 y = 0.51+0.55x P = 0.43
R = 0.13 R2adj = −0.028 y = 0.67+0.39x P = 0.55
H
Stem Cell
Lipid Metabolism
Immunomodulatory
ISC CD HFD
4kb 3kb 2kb 2kb 3kb 2kb 1kb 3kb 2kb
[0-2.05] [0-1.02] [0-1.15] [0-0.2] [0-1] [0-0.67] [0-0.45] [0 - 0.29] [0-0.15]
Olfm4 Ascl2 Axin2 Fabp1 Acaa1b Cpt2 Gbp10 Nlrp9b Mir29b-2
P = 0.716 P = 0.229 P = 0.515 P = 2.84e-3 P = 7.17e-4 P = 1.02e-4 P = 0.013 P = 0.018 P = 0.0301
Normalized Counts
CD HFD

### Figure S2

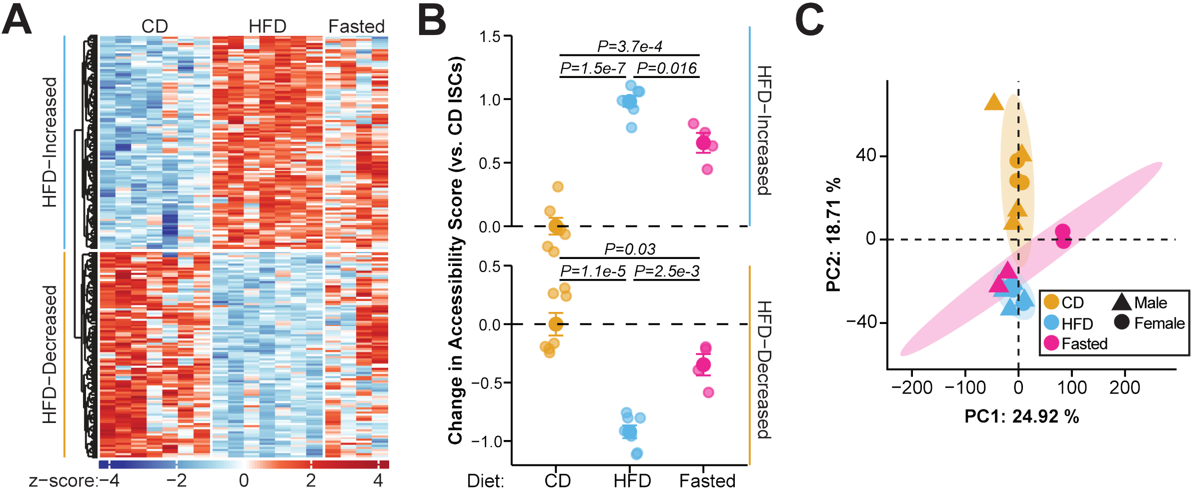
A
CD
HFD
Fasted
HFD−Increased
HFD−Decreased
z−score: −4 −2 0 2 4
B
Change in Accessibility Score (vs. CD ISCs)
P=3.7e-4
P=1.5e-7
P=0.016
P=0.03
P=1.1e-5
P=2.5e-3
HFD−Increased
HFD−Decreased
Diet: CD HFD Fasted
C
PC1: 24.92 %
PC2: 18.71 %
CD
HFD
Fasted
Male
Female

### Figure S3

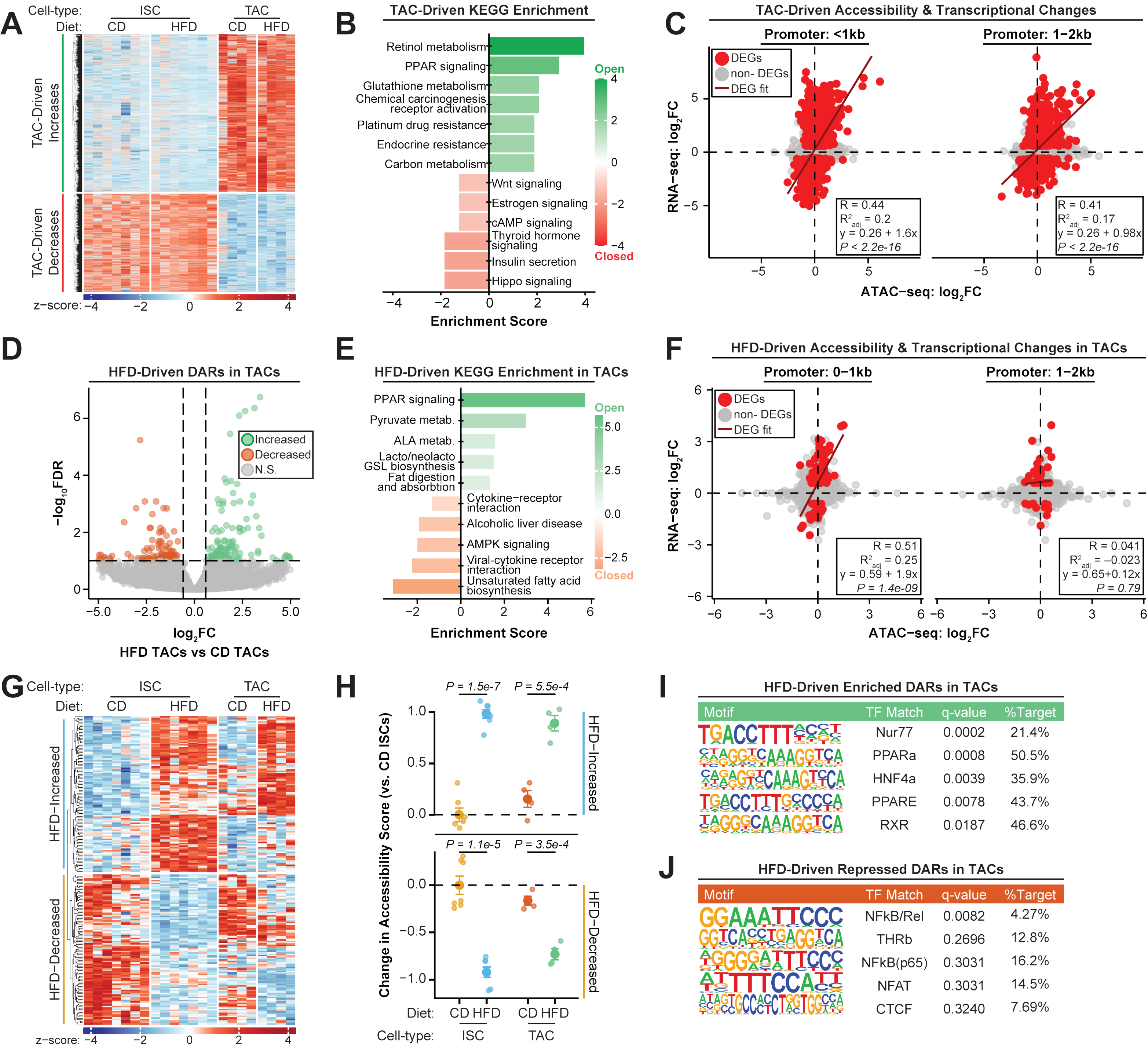
A
Cell-type: ISC TAC
Diet: CD HFD CD HFD
TAC-Driven Increases
TAC-Driven Decreases
z-score: −4 −2 0 2 4
B
TAC-Driven KEGG Enrichment
Retinol metabolism
PPAR signaling
Glutathione metabolism
Chemical carcinogenesis receptor activation
Platinum drug resistance
Endocrine resistance
Carbon metabolism
Wnt signaling
Estrogen signaling
cAMP signaling
Thyroid hormone signaling
Insulin secretion
Hippo signaling
Open
Closed
Enrichment Score
C
TAC-Driven Accessibility & Transcriptional Changes
Promoter: <1kb
Promoter: 1−2kb
DEGs
non- DEGs
DEG fit
R = 0.44
R²adj = 0.2
y = 0.26 + 1.6x
P < 2.2e-16
R = 0.41
R²adj = 0.17
y = 0.26 + 0.98x
P < 2.2e-16
RNA−seq: log2FC
ATAC−seq: log2FC
D
HFD-Driven DARs in TACs
Increased
Decreased
N.S.
−log10FDR
log2FC
HFD TACs vs CD TACs
E
HFD-Driven KEGG Enrichment in TACs
PPAR signaling
Pyruvate metab.
ALA metab.
Lacto/neolacto GSL biosynthesis
Fat digestion and absorbtion
Cytokine−receptor interaction
Alcoholic liver disease
AMPK signaling
Viral-cytokine receptor interaction
Unsaturated fatty acid biosynthesis
Open
Closed
Enrichment Score
F
HFD-Driven Accessibility & Transcriptional Changes in TACs
Promoter: 0−1kb
Promoter: 1−2kb
DEGs
non- DEGs
DEG fit
R = 0.51
R²adj = 0.25
y = 0.59 + 1.9x
P = 1.4e-09
R = 0.041
R²adj = −0.023
y = 0.65+0.12x
P = 0.79
RNA−seq: log2FC
ATAC−seq: log2FC
G
Cell-type: ISC TAC
Diet: CD HFD CD HFD
HFD−Increased
HFD−Decreased
z-score: −4 −2 0 2 4
H
P = 1.5e-7
P = 5.5e-4
P = 1.1e-5
P = 3.5e-4
HFD−Increased
HFD−Decreased
Change in Accessibility Score (vs. CD ISCs)
Diet: CD HFD CD HFD
Cell-type: ISC TAC
I
HFD-Driven Enriched DARs in TACs
Motif | TF Match | q-value | %Target
Nur77 | 0.0002 | 21.4%
PPARa | 0.0008 | 50.5%
HNF4a | 0.0039 | 35.9%
PPARE | 0.0078 | 43.7%
RXR | 0.0187 | 46.6%
J
HFD-Driven Repressed DARs in TACs
Motif | TF Match | q-value | %Target
NFkB/Rel | 0.0082 | 4.27%
THRb | 0.2696 | 12.8%
NFkB(p65) | 0.3031 | 16.2%
NFAT | 0.3031 | 14.5%
CTCF | 0.3240 | 7.69%

### Figure S4

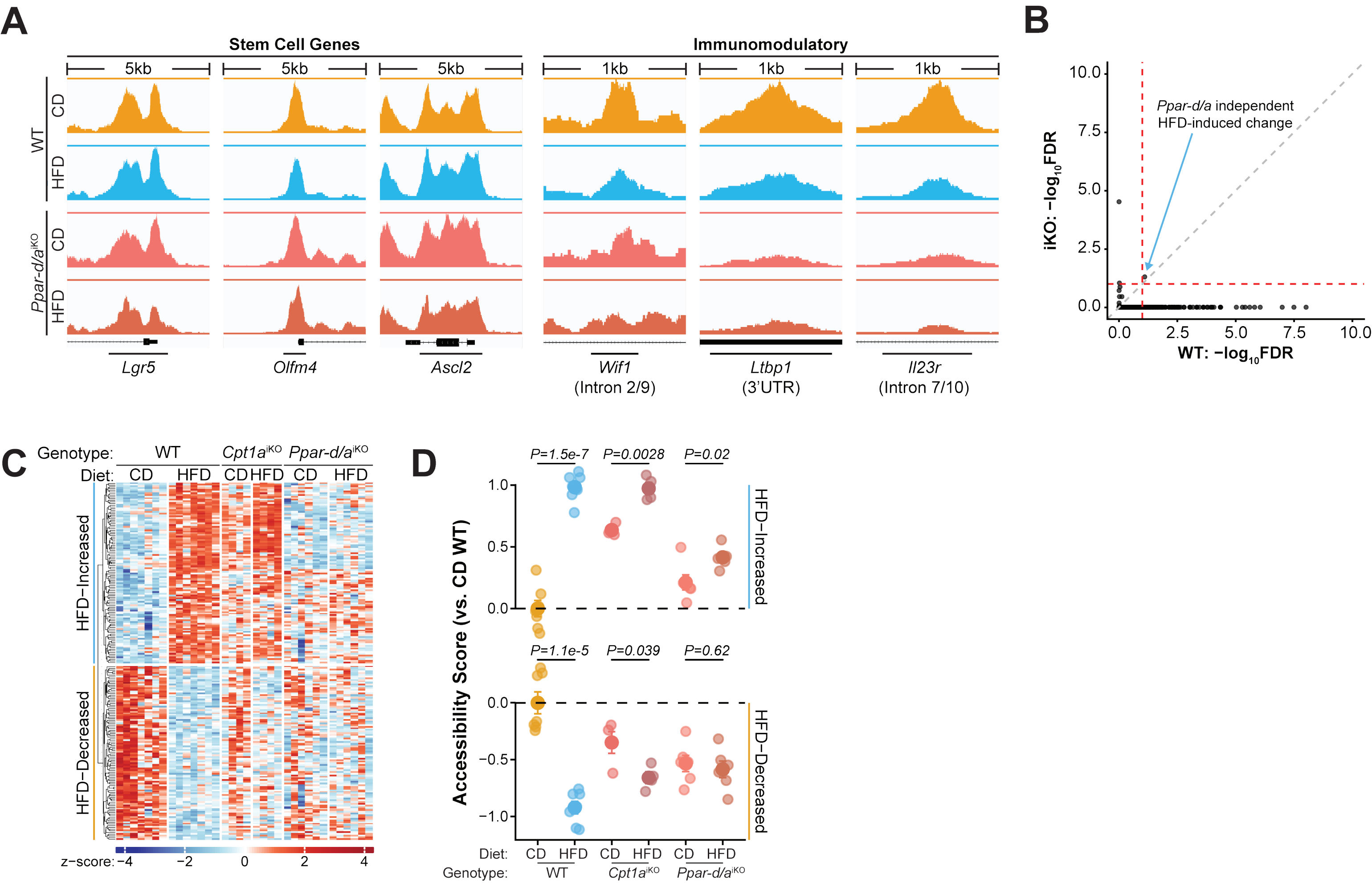
A
Stem Cell Genes
Immunomodulatory
5kb
1kb
WT
CD
HFD
Ppar-d/a iKO
Lgr5
Olfm4
Ascl2
Wif1 (Intron 2/9)
Ltbp1 (3'UTR)
Il23r (Intron 7/10)
B
Ppar-d/a independent HFD-induced change
iKO: −log10FDR
WT: −log10FDR
C
Genotype: WT Cpt1a iKO Ppar-d/a iKO
Diet: CD HFD CD HFD CD HFD
HFD−Increased
HFD−Decreased
z−score: −4 −2 0 2 4
D
P=1.5e-7
P=0.0028
P=0.02
P=1.1e-5
P=0.039
P=0.62
Accessibility Score (vs. CD WT)
HFD−Increased
HFD−Decreased
Diet: CD HFD CD HFD CD HFD
Genotype: WT Cpt1a iKO Ppar-d/a iKO

### Figure S5

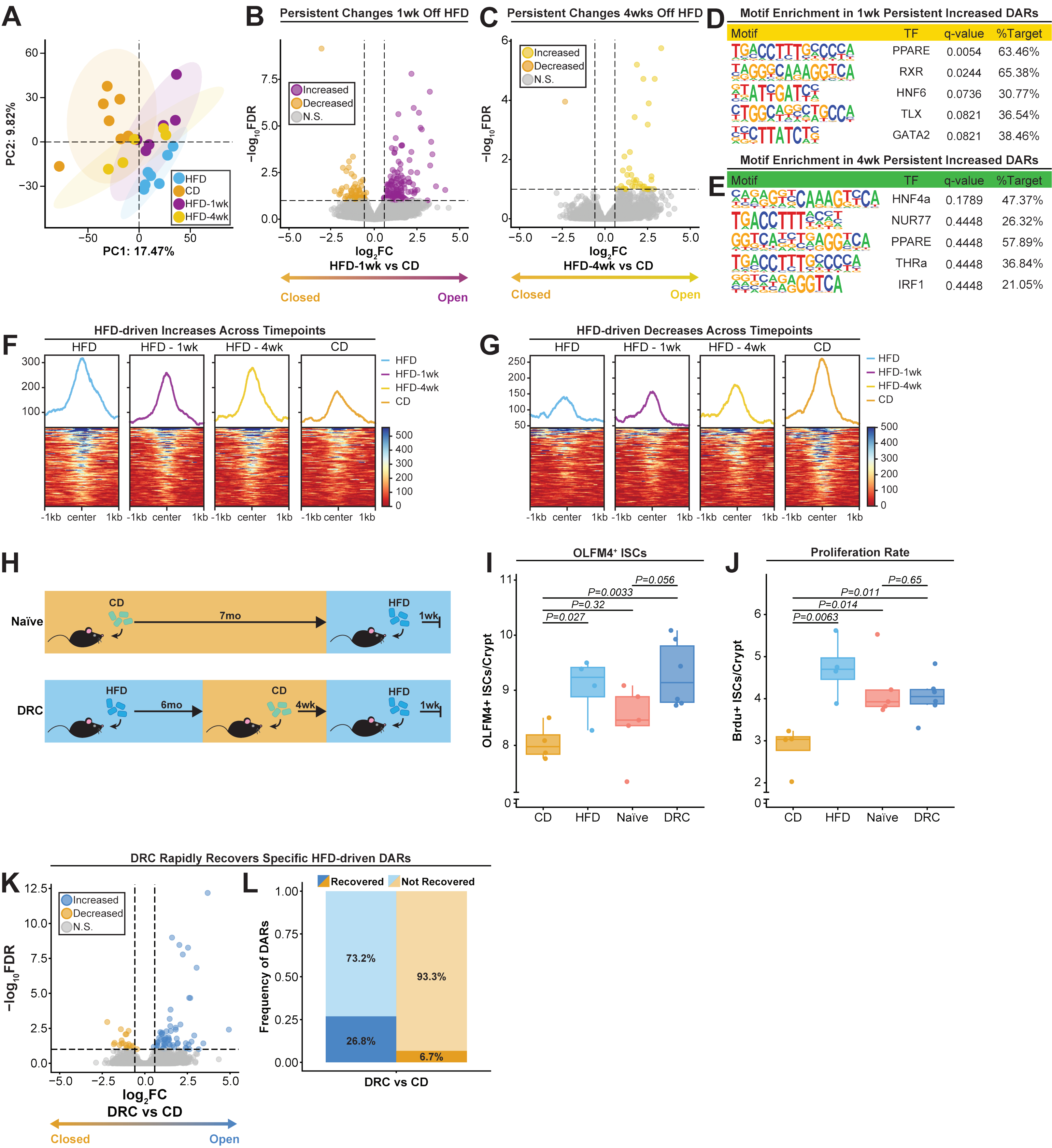
A
PC2: 9.82%
PC1: 17.47%
HFD
CD
HFD-1wk
HFD-4wk
B
Persistent Changes 1wk Off HFD
Increased
Decreased
N.S.
−log10FDR
log2FC
HFD-1wk vs CD
Closed
Open
C
Persistent Changes 4wks Off HFD
Increased
Decreased
N.S.
log2FC
HFD-4wk vs CD
Closed
Open
D
Motif Enrichment in 1wk Persistent Increased DARs
Motif TF q-value %Target
PPARE 0.0054 63.46%
RXR 0.0244 65.38%
HNF6 0.0736 30.77%
TLX 0.0821 36.54%
GATA2 0.0821 38.46%
E
Motif Enrichment in 4wk Persistent Increased DARs
Motif TF q-value %Target
HNF4a 0.1789 47.37%
NUR77 0.4448 26.32%
PPARE 0.4448 57.89%
THRa 0.4448 36.84%
IRF1 0.4448 21.05%
F
HFD-driven Increases Across Timepoints
HFD
HFD - 1wk
HFD - 4wk
CD
-1kb center 1kb
G
HFD-driven Decreases Across Timepoints
HFD
HFD - 1wk
HFD - 4wk
CD
-1kb center 1kb
H
Naïve
CD
7mo
HFD
1wk
DRC
HFD
6mo
CD
4wk
HFD
1wk
I
OLFM4+ ISCs
OLFM4+ ISCs/Crypt
P=0.056
P=0.0033
P=0.32
P=0.027
CD HFD Naïve DRC
J
Proliferation Rate
Brdu+ ISCs/Crypt
P=0.65
P=0.011
P=0.014
P=0.0063
CD HFD Naïve DRC
K
DRC Rapidly Recovers Specific HFD-driven DARs
Increased
Decreased
N.S.
−log10FDR
log2FC
DRC vs CD
Closed
Open
L
Recovered
Not Recovered
Frequency of DARs
73.2%
26.8%
93.3%
6.7%
DRC vs CD

### Figure S6

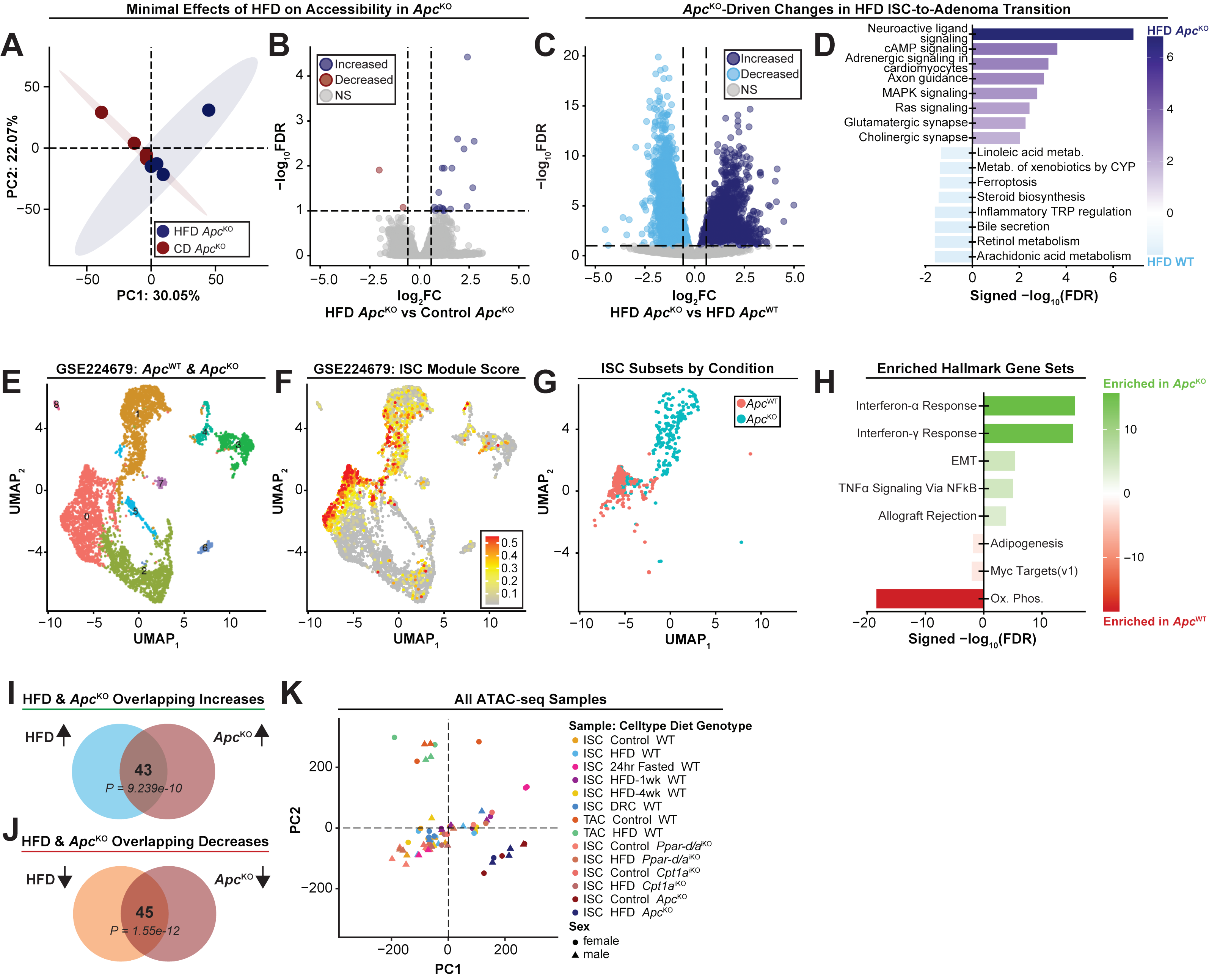
Minimal Effects of HFD on Accessibility in Apc^KO
A
PC1: 30.05%
PC2: 22.07%
HFD Apc^KO
CD Apc^KO
B
Increased
Decreased
NS
log2FC
−log10FDR
HFD Apc^KO vs Control Apc^KO
Apc^KO-Driven Changes in HFD ISC-to-Adenoma Transition
C
Increased
Decreased
NS
log2FC
−log10FDR
HFD Apc^KO vs HFD Apc^WT
D
Neuroactive ligand signaling
cAMP signaling
Adrenergic signaling in cardiomyocytes
Axon guidance
MAPK signaling
Ras signaling
Glutamatergic synapse
Cholinergic synapse
Linoleic acid metab.
Metab. of xenobiotics by CYP
Ferroptosis
Steroid biosynthesis
Inflammatory TRP regulation
Bile secretion
Retinol metabolism
Arachidonic acid metabolism
HFD Apc^KO
HFD WT
Signed −log10(FDR)
E
GSE224679: Apc^WT & Apc^KO
UMAP1
UMAP2
F
GSE224679: ISC Module Score
G
ISC Subsets by Condition
Apc^WT
Apc^KO
H
Enriched Hallmark Gene Sets
Interferon-α Response
Interferon-γ Response
EMT
TNFα Signaling Via NFkB
Allograft Rejection
Adipogenesis
Myc Targets(v1)
Ox. Phos.
Enriched in Apc^KO
Enriched in Apc^WT
Signed −log10(FDR)
I
HFD & Apc^KO Overlapping Increases
HFD
Apc^KO
43
P = 9.239e-10
J
HFD & Apc^KO Overlapping Decreases
HFD
Apc^KO
45
P = 1.55e-12
K
All ATAC-seq Samples
PC1
PC2
Sample: Celltype Diet Genotype
ISC Control WT
ISC HFD WT
ISC 24hr Fasted WT
ISC HFD-1wk WT
ISC HFD-4wk WT
ISC DRC WT
TAC Control WT
TAC HFD WT
ISC Control Ppar-d/a^iKO
ISC HFD Ppar-d/a^iKO
ISC Control Cpt1a^iKO
ISC HFD Cpt1a^iKO
ISC Control Apc^KO
ISC HFD Apc^KO
Sex
female
male
